# ssJSD: A Fusion of Sparsity and Spatial Information for HiC Single-Cell Clustering

**DOI:** 10.64898/2026.09.14.751457

**Authors:** Sang Wan Lee, Shili Lin

**Author notes:** Address for correspondence: Shili Lin, PhD, Department of Statistics The Ohio State University 1958 Neil Avenue, Columbus, OH 43210-1247, USA.

## Abstract

Single-cell high-throughput chromatin conformation capture (scHiC) enables profiling three-dimensional genome architecture at cellular resolution, providing insights into cell-to-cell variability and cellular functions. Recent frameworks utilize spatial interaction patterns to derive dissimilarity measures for downstream tasks such as cell clustering. However, the inherent sparsity and ultra-high dimensionality of scHiC contact matrices pose significant challenges. A central hurdle is that existing measures typically treat all zeros without distinction, failing to differentiate biologically meaningful structural zeros (SZs) from technical dropouts. Here, we introduce ssJSD (spatial and sparsity informed Jensen-Shannon Divergence), a computational framework designed to explicitly account for scHiC-specific sparsity patterns. By integrating band-wise contact frequency profiles with SZ-induced sparsity matrices, ssJSD leverages both spatial interaction patterns and biological absence of contacts. We adopted two complementary integration strategies: an early fusion approach that concatenates information into a single representation, and a late fusion approach that integrates JSD-based dissimilarities through diverse averaging methods. Through simulations and applications to human cell lines and prefrontal cortex data, we demonstrate that ssJSD improves clustering accuracy and effectively distinguishes cell types. Our study indicates that integrating SZ patterns is important for accurately quantifying cell-to-cell variability in 3D genomics.

## 1 Introduction

The human genome is organized into a complex three-dimensional (3D) structure that plays a fundamental role in gene regulation and cellular function [1, 2, 3]. Recent advances in singlecell genomics have provided unprecedented opportunities to investigate the 3D structure at the level of individual cells, revealing substantial cell-to-cell variability in chromatin organization [4, 5, 6, 7]. Single-cell HiC (scHiC) is one such technology that captures genome-wide chromatin contacts within individual cells, opening the possibility for identifying cellular sub-populations based on their 3D genome architecture [4, 8, 9, 10]. However, extracting such insights from scHiC data remains technically challenging due to both their unique structure and extreme sparsity.

A defining feature of scHiC data is their representation as two-dimensional (2D) contact matrices, where each entry reflects the observed interaction frequency between a pair of genomic segments (referred to as a locus pair, or LP). A well-established property of these matrices is that contact frequency decays systematically with increasing one-dimensional (1D) genomic distance [1]. As a result, the matrices display spatial patterns along genomic distance bands, with LPs having similar 1D distance sharing similar interaction profiles. This distance-dependent structure motivated the development of stratum-based approaches such as HiCRep [11] and JSD [12], which aggregate 2D contact information into band-wise summaries for downstream analyses.

Beyond the spatial structure of contact matrices, scHiC data suffer from extreme sparsity, with zero entries often comprising more than 90% of all observations [13, 14, 15]. Each cell produces a matrix with millions of LPs, yet interactions among a vast majority are unobserved [16, 17]. Several strategies have been proposed to mitigate the effects of sparsity: smoothing-based methods such as SCL [18] and GenomeDISCO [19] recover signal by averaging over neighboring entries, while graph-based and deep learning frameworks such as scHiCluster [13] and Higashi [20] construct embeddings that are robust to sparse observations.

Nevertheless, a central challenge—not all zeros in scHiC are equivalent—has not been sufficiently addressed. Among the observed zeros (OZs), some represent true biological absence of interaction, known as structural zeros (SZs), while others arise from technical factors, known as dropouts (DOs) [14, 21, 22]. Existing methods for computing cell-to-cell dissimilarity, whether based on stratum-based measure [11, 12] or low-dimensional embeddings [13, 20], typically treat all zeros uniformly, ignoring this distinction. HiCImpute [21] and scHiCSRS [22] explicitly model the SZ/DO distinction using probabilistic frameworks. Apart from a recent method, szKendall [23], SZ information has not been explicitly used to calculate dissimilarity measures used for clustering.

From szKendall, we observe dramatic improvements in the power for detecting structural differences among cells; however, szKendall is computationally expensive. To address the computational intensity while leveraging the valuable information contained in SZs, we propose spatial and sparsity-informed Jensen-Shannon divergence (ssJSD), which combines the stratum-based JSD framework [12] with SZ annotations for computational efficiency. Rather than treating all OZs uniformly, ssJSD leverages probabilistic SZ predictions from imputation models to construct two representations of each cell that are then integrated using two strategies. Under the early fusion strategy, the contact and SZ information are concatenated into a unified cell representation before computing JSD. Under the late fusion strategy, JSD is computed separately from each representation and the resulting dissimilarities are combined. This design allows ssJSD to capture both contact frequency variation and SZ-driven structural differences between cells, facilitating more precise quantification of chromatin architecture variability across cells.

We assess the performance of ssJSD through simulation studies designed to reflect the range of conditions encountered in real scHiC experiments, including varying levels of sparsity, sequencing depth, and degree of separation between cell types. We additionally apply ssJSD to two real datasets, comprising five human cell lines and human prefrontal cortex data. We demonstrate that ssJSD substantially improves clustering performance compared to traditional approaches.

## 2 Results

### 2.1 Overview of ssJSD

We devised **ssJSD** to improve cell clustering performance by jointly utilizing the SZ-induced sparsity matrix and the contact frequency matrix from HiC single-cell contact frequencies.

HiC data are organized into a 2D matrix of contact frequencies containing *spatial* information. Specifically, contacts between LPs that are closer in 1D genomic distance typically occur more frequently (i.e., having a larger contact frequency, or CF) [1], unless such interactions are prevented by underlying biological mechanisms. Since LPs on a band parallel to the main diagonal of the matrix share the same genomic distance, aggregating along these diagonal bands preserves *spatial* information while also mitigating, to some extent, the sparsity problem in single-cell data [11, 12]. This leads to our condensed vector representation of the original 2D contact frequency matrix, *P_CF_* = (*C*_1_*, C*_2_*, . . . , C_B_*), where *C*_1_ is the total contact frequency along the band one off the main diagonal, and so on, and *B* is the number of bands, one less than the size of the 2D matrix (Figure 1a).

**Figure 1:**
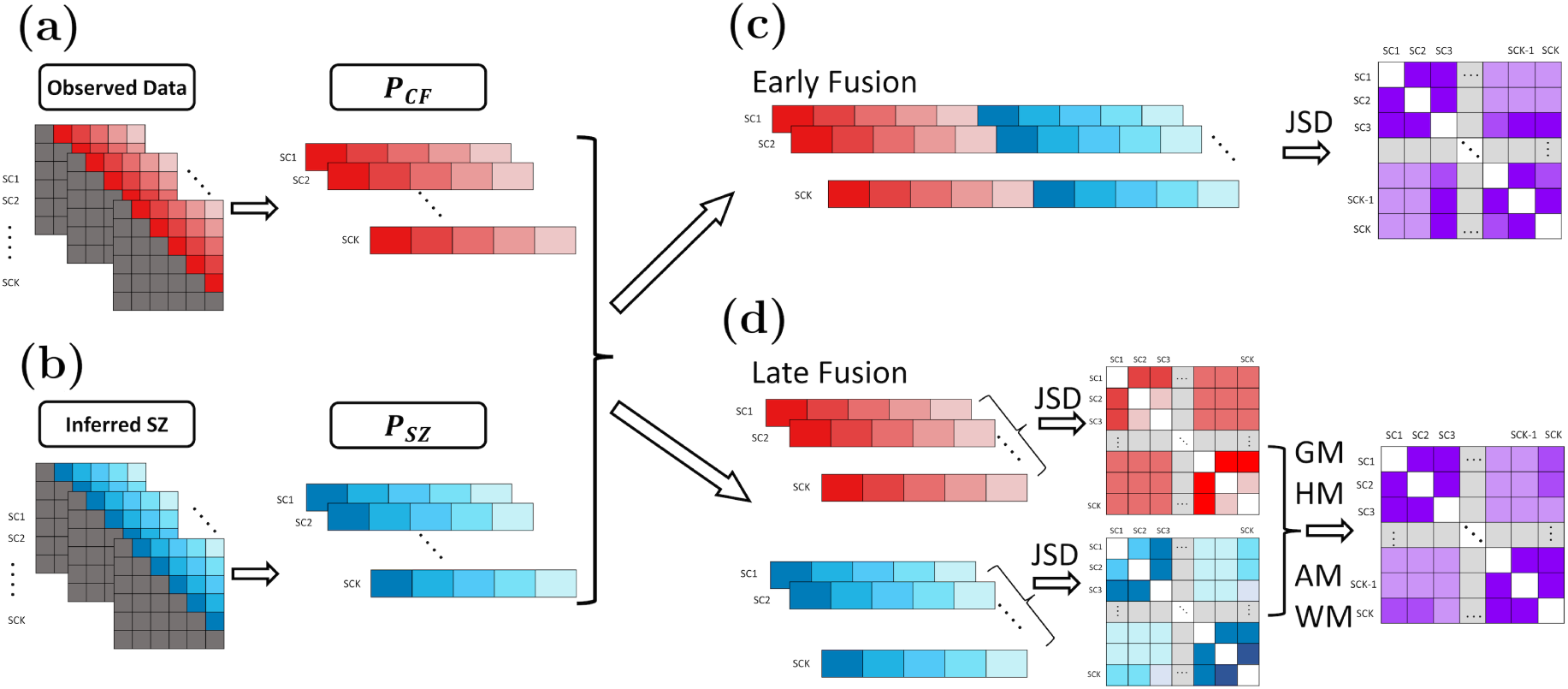
Overview of ssJSD. (a) Observed HiC 2D contact matrices of single cells, with each same-colored entry representing a band parallel to the main diagonal, which are aggregated to form the *P_CF_* probability vector for each cell after appropriate normalization. (b) Inferred SZ matrices with binary values indicating whether an observed zero is an SZ or a DO, which are then aggregated to form the *P_SZ_* probability vector for each cell after appropriate normalization. (c) An early-fusion framework, which first concatenates the two probability vectors from the contact matrix and inferred SZ matrix, then computes the JSD for each pair of the re-normalized concatenated probability distributions. (d) A late-fusion framework, in which a JSD is computed for each pair of probability distributions for *P_CF_* and *P_SZ_* separately, then the two dissimilarity matrices are combined using one of four variants: geometric mean (GM); harmonic mean (HM); arithmetic mean (AM); weighted mean (WM).

Although aggregating data along diagonal bands helps address sparsity to some extent, not all zero contact frequencies are meaningless. In fact, SZ-induced sparsity may contain rich information about cell lineage, and leveraging such information can provide valuable signals for cell clustering. To this end, we construct another data matrix—the SZ-informed sparsity matrix—a binary matrix with 0/1 entries, where 1 denotes that the two loci do not interact (i.e., a structural zero). This matrix can be inferred using existing methods [21, 22]. Analogous to the contact frequency matrix, we also build a vector representation of this matrix, *P_SZ_* (Figure 1b). This vector reflects both *spatial* and *structure-zero-aware* information through SZs.

We then propose two fusion approaches to combine the information from these two vectors to capture dissimilarities among a set of cells. The first approach is referred to as *early fusion*, where the two vectors are merged at the first step, after normalizing each by twice the sum of its entries. We then compute the JSD across these probability vectors to form a dissimilarity matrix (Figure 1c). The second approach is referred to as *late fusion*, where two JSD matrices are computed separately, one from each vector, and then combined using four different averaging methods: geometric mean (GM), harmonic mean (HM), arithmetic mean (AM), and weighted mean (WM) (Figure 1d).

By construction, ssJSD leverages the additional information contained in SZ patterns, which reflect the true absence of chromatin interactions. Our approach provides an improved representation of both the spatial organization of chromosomal contacts and the cell-to-cell variability arising from SZ patterns.

### 2.2 ssJSD separates cell clusters by utilizing SZ information

We generated HiC data under six scenarios of sparsity and sequencing depth (Table 1; Sim1–Sim6) while preserving spatial information based on a K562 scHiC dataset [9]. Each dataset consists of 150 single-cell contact matrices, along with information on whether an observed zero is a structural zero or a dropout. These 150 single cells are evenly divided into three “subtypes” using two designs. In Design 1, the main difference between the subtypes is due to different sparsity profiles, as the underlying 3D structure of a single K562 cell was used (accession: GSM2109974). In Design 2, the differences between the three subtypes are due to both sparsity profiles and 3D structures, since the contact matrices for each subtype were generated based on the underlying 3D structures of three different K562 cells (GSM2109974, GSM2109985, and GSM2109993). For each subtype, 50 single-cell contact matrices were generated following a three-step procedure (Methods).

**Table 1:**
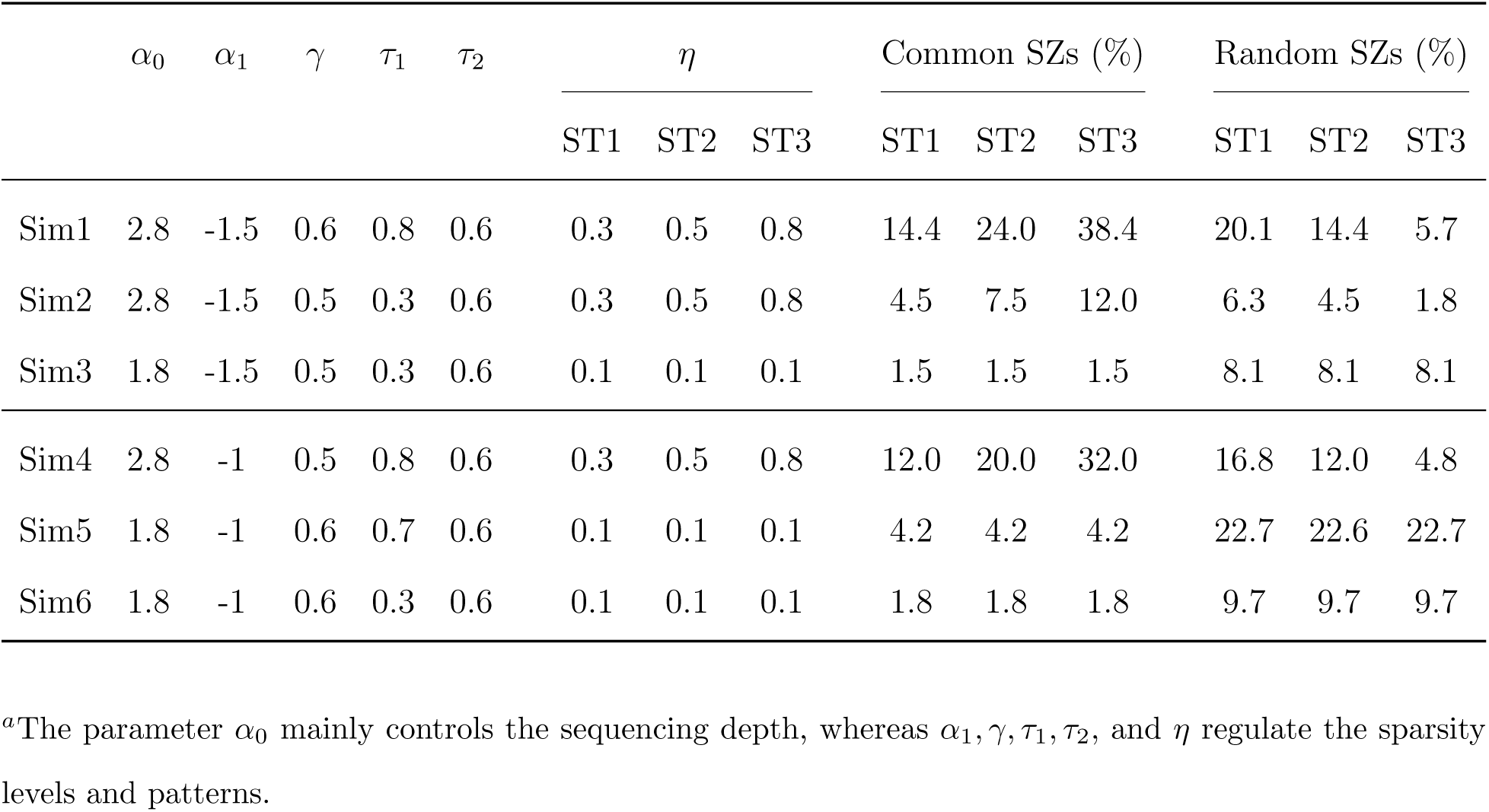
Specification of parameters*^a^* and resulting percentages of two types of structural zeros.

The two designs pairing with each of the six sparsity and sequencing depth scenarios yield a total of 12 settings (Design 1, Sim1–Sim6; Design 2, Sim1–Sim6), providing a rich resource for a comprehensive evaluation of the novel ssJSD methodology as well as existing methodologies based on the original JSD. Specifically, for ssJSD, we evaluated the early fusion (ssJSD-EF) and four versions of late fusion (ssJSD-AM, ssJSD-GM, ssJSD-HM, ssJSD-WM). To highlight the contribution from the sparsity profile, we also considered two additional variants—ssJSD-AM(OZ) and ssJSD-GM(OZ)—which are counterparts of ssJSD-AM and ssJSD-GM, except that the sparsity matrix is constructed directly from observed zeros (i.e., all observed zeros are erroneously treated as structural zeros). For the original JSD methodology [12], we applied three versions of the data: the observed contact matrices (JSD-OBS), the Gaussian kernel–improved contact matrices (JSD-GK), and the random walk–improved contact matrices (JSD-RW), where the GK- and RW-improved data were obtained based on algorithms from the existing literature [18, 19]. In total, we compared 10 methods.

For the six settings in which the differences between the three subtypes are due only to sparsity (Design 1, Sim1–Sim6), it is not surprising to see that none of the three versions of the original JSD (i.e. JSD-OBS, JSD-GK, and JSD-RW) was able to delineate the subtypes. The same was true when sparsity matrices were constructed solely from observed zeros (ssJSD-AM(OZ) and ssJSD-GM(OZ)). Specifically, the heatmap representations of the dissimilarity matrices and three commonly used 2D projections— UMAP [24], t-SNE [25], and MDS [26]— all show the mixing of cells from the three subtypes (Supplementary Figures S1-S6, top panels). In contrast, when the true sparsity matrices were used, the three subtypes were well separated, either using the early fusion method (ssJSD-EF) or the four late fusion methods (ssJSD-AM, ssJSD-GM, ssJSD-HM, ssJSD-WM). This is evident based on all evaluation metrics: the heatmaps and the 2D projections (Supplementary Figures S1-S6, bottom panels).

For the six settings in which the differences between the three subtypes are due to both sparsity profiles and spatial structures (Design 2, Sim1–Sim6), JSD and ssJSD methods with erroneous SZ sparsity information performed well for the Sim1, Sim2, and Sim4 datasets—where sequencing depth was high—with the exception of JSD-RW (Supplementary Figures S7-S9, top panels). However, with decreased sequencing depth, relying solely on spatial structural differences was no longer sufficient: the three subtypes became entangled in both the heatmap visualizations and the three 2D projections (Figure 2 for Sim3 and Supplementary Figures S10-S11 for Sim5 and Sim6, top panels). In contrast, by incorporating SZ sparsity information, all five ssJSD methods successfully separated the three subtypes across all six scenarios (Figure 2, Supplementary Figures S7-S11, bottom panels).

**Figure 2:**
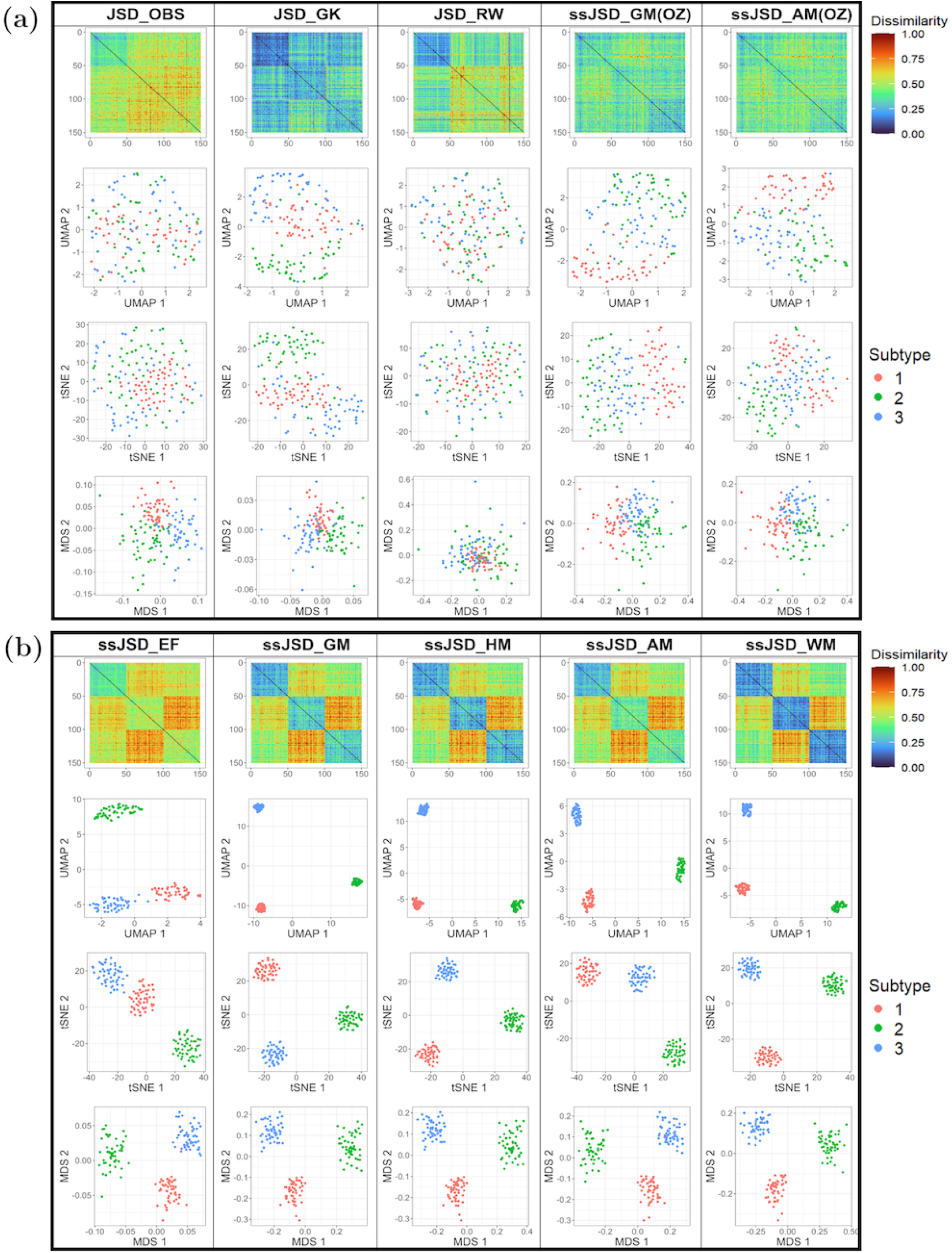
Comparison of ten methods in terms of heatmap and 2D projections of the dissimilarity matrix for Design 2 Sim3. (a) Five methods that do not use correct SZ information; (b) Five proposed methods that use SZ information.

### 2.3 ssJSD consistently achieves excellent performance under four evaluation criteria

The performance of the original JSD and the novel ssJSD methodologies was further evaluated for performance variability using 100 simulated datasets for each of the 12 settings. Clustering was performed either directly on dissimilarity matrices using Partitioning Around Medoids (PAM) [27] or on three 2D projection results using K-means (KM) [28]. The clustering performance was assessed with four criteria: Adjusted Rand Index (ARI), the proportion of within-cluster sum of squares to the total sum of squares (WSS), Average Silhouette Width (ASW), and Minimum Isolation Score (MIS) (Methods). In this evaluation, we also included four counterparts of the ssJSD late fusion variants, with all OZs treated as SZs.

Consistent with the results from one simulated dataset per scenario for Design 1, where the differences among the three subtypes are mainly driven by sparsity, none of the methods that relied solely on spatial structure information from the contact matrix performed well—this set included the ssJSD methods late fusion versions with all OZ treated as SZs.

These methods yielded low ARI, ASW, and MIS, but high WSS. This observation held regardless of whether clustering was performed directly on the dissimilarity matrix or on the 2D projection matrix across all six scenarios (Figure 3, top panel for showing the average ARI, and Supplementary Figures S12-S17 displaying variability across the 100 datasets for all four criteria).

**Figure 3:**
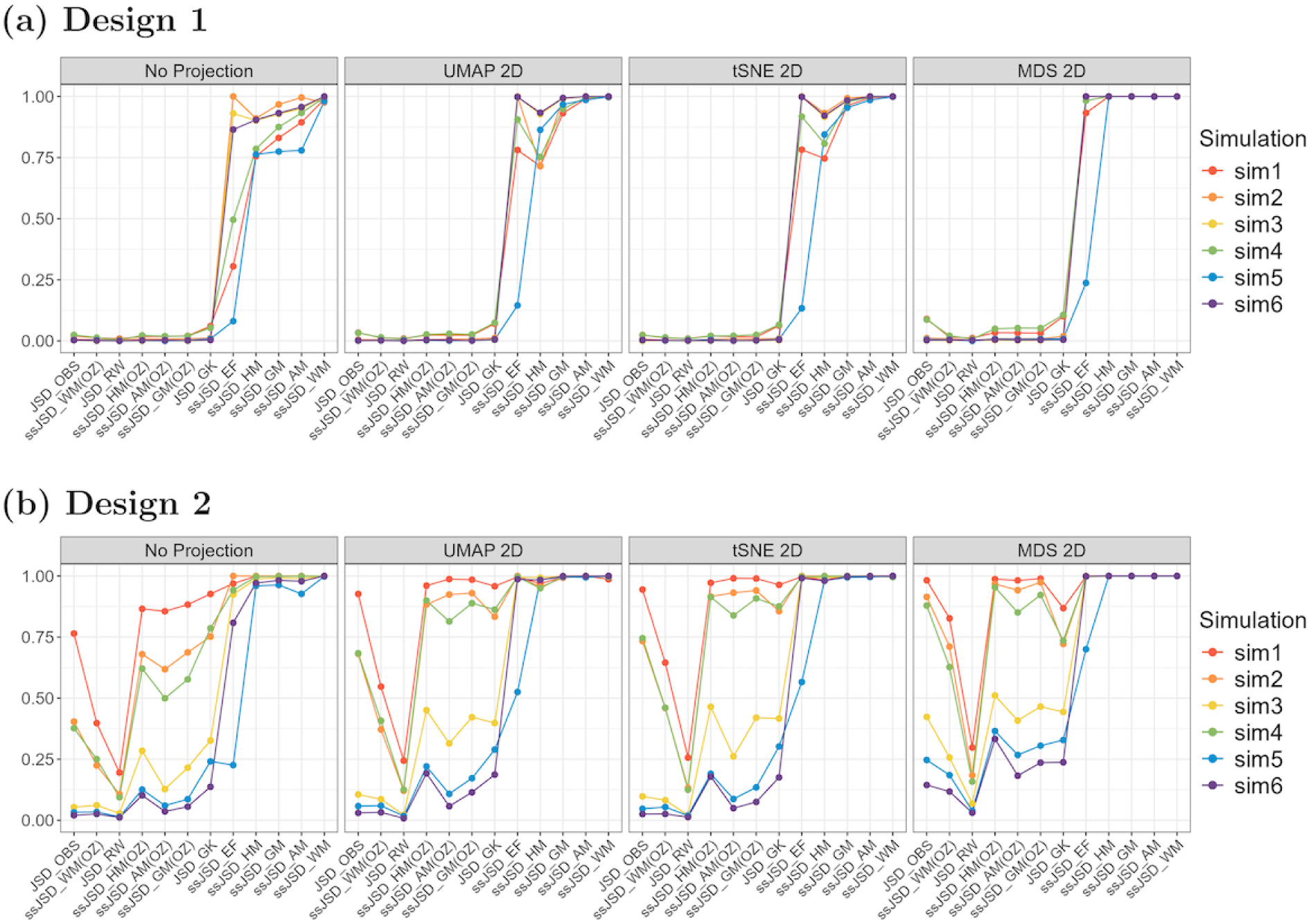
Comparison of the average ARI for 12 methods based on the dissimilarity matrix (no projection), or three methods of 2D projections: UMAP, tSNE, and MDS. (a) Results for all six sparsity and sequencing depth scenarios under Design 1; (b) Results for all six sparsity and sequencing depth scenarios under Design 2. For both the (a) and (b) panels, the methods were ordered based on the performance (from worst to best) for the Design 1, Sim1 data.

In contrast, all five ssJSD methods that incorporated sparsity information from structural zeros consistently produced higher ARI and ASW, and lower WSS, across the board, regardless of whether dimensionality reduction was applied. For MIS, we observed a substantial improvement for clustering results based on UMAP and t-SNE projections, especially for the ssJSD-AM and ssJSD-WM methods (Supplementary Figures S12-S17).

For Design 2, our numerical evaluation of clustering results also agrees with the visualization results presented in the previous subsection. For datasets with sufficient sequencing depth (Sim1, Sim2, and Sim4), most methods relying only on contact count matrices performed reasonably well, with the exception of JSD-RW (random-walk-smoothed data) (Figure 3 bottom panel and Supplementary Figures S18-S23). When sequencing depth was reduced (Sim3, Sim5, and Sim6), methods using only contact count matrices were no longer able to separate the clusters. Consistent with the Design 1 results, all five ssJSD variants yielded high ARI and ASW values, low WSS, and clearly outperformed methods that did not use SZ-induced sparsity information. As in the previous subsection, UMAP and t-SNE projections continued to produce tight clusters, reflected in the MIS values being greater than 1.

### 2.4 ssJSD still outperforms under a range of SZ mis-specifications

As shown in the previous two subsections, ssJSD, by further utilizing the SZ-induced sparsity matrix in addition to the contact count matrix, substantially enhanced clustering accuracy, producing well-defined and clearly separated subtypes under two designs on spatial structures. Nevertheless, those findings were based on the idealized assumption that all SZs were perfectly detected. In reality, achieving flawless identification of true SZs is rarely feasible. To overcome this limitation, we examined the robustness of ssJSD under different levels of sensitivity (the fraction of true SZs correctly recognized) and specificity (the fraction of true DOs correctly classified).

For each of the 12 simulation settings and 100 datasets under each setting, we evaluated 42 distinct sensitivity–specificity mis-specifications. In the first 21 cases, sensitivity was fixed at *α* = 0.95, 0.8, or 0.6 by randomly reassigning 1 *− α* proportion of true SZs as DOs, while specificity was varied from *β* = 0.9 down to 0.3 in decrements of 0.1 by relabeling 1 *– β* proportion of true DOs as SZs. In the other 21 cases, the roles of sensitivity and specificity were reversed, with specificity fixed while sensitivity varying in a range.

We applied the PAM clustering algorithm to all 100 datasets for each combination of the 12 simulation settings and the 42 specificity-sensitivity specifications. Examining the mean ARI patterns across the 100 datasets, we observe that for Design 1—where subtype separation relies more on the SZ-induced sparsity matrix—the ARI declines rapidly as either sensitivity or specificity decreases. Nevertheless, a respectable ARI of around 0.7 can still be achieved when both sensitivity and specificity are reasonably high (e.g., at least 0.8). More importantly, even with a low sensitivity or specificity (e.g., sensitivity = 0.6 and specificity = 0.3, or vice versa), the ARI for any of the late fusion ssJSD methods rarely falls below the baseline defined by JSD-OBS (Figure 4, top panel, for Sim2, and Supplementary Figures S24-S28, top panels, for the other five sparsity-sequencing depth scenarios).

**Figure 4:**
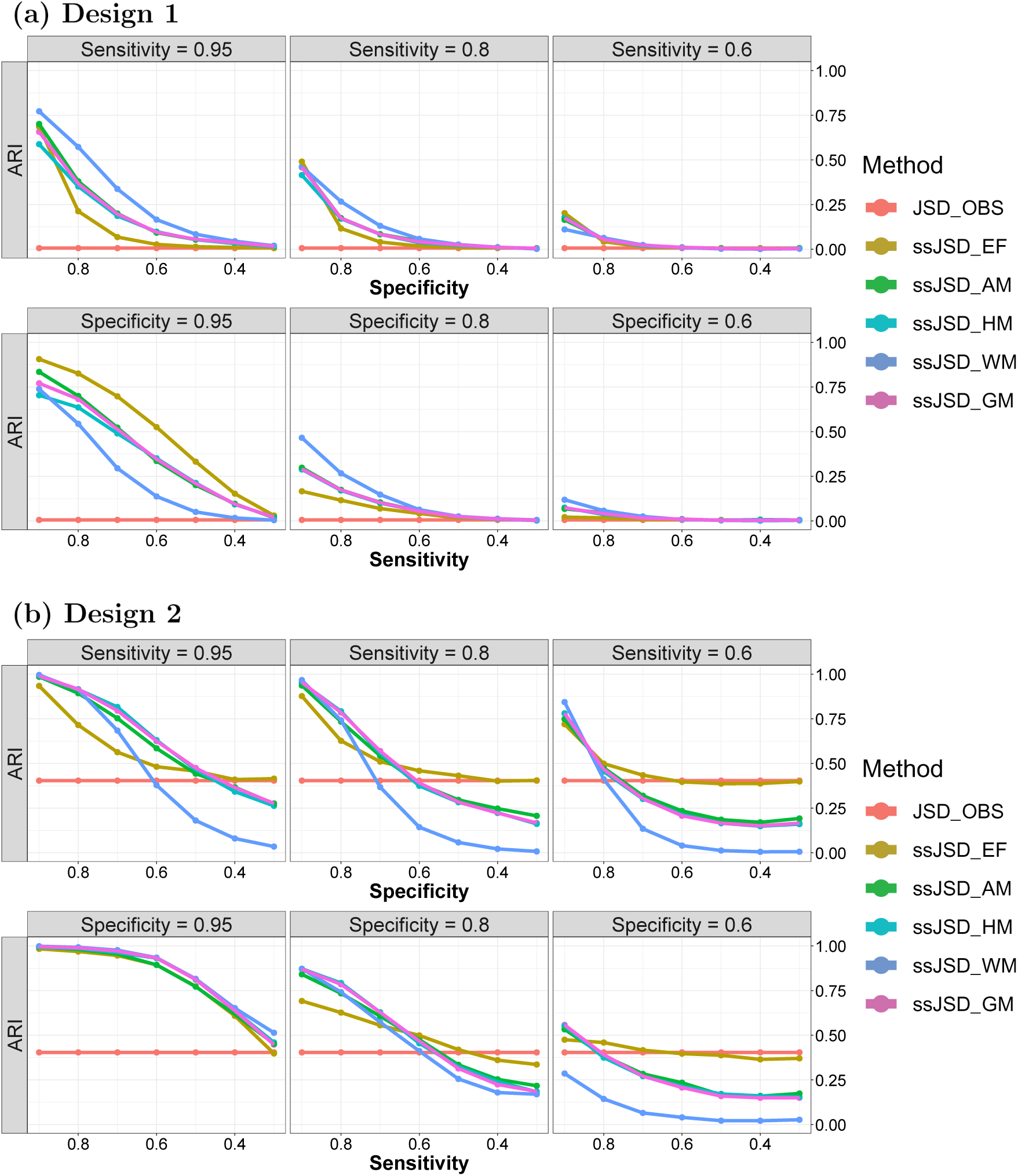
Effect of specificity and sensitivity on clustering performance based on ARI. (a) Design 1 Sim2 data, where each plot in the first row has a fixed sensitivity but varying specificity from 0.9 to 0.3 with a decrement of 0.1, and each plot in the second row has a fixed specificity but varying sensitivity from 0.9 to 0.3 with a decrement of 0.1; (b) Same as (a) but for Design 2 Sim2 data.

For data generated under Design 2 with high sequencing depth (Sim1, Sim2, and Sim4), the baseline ARIs from JSD-OBS are already high; however, nearly all ssJSD methods outperform the baseline when either sensitivity or specificity is reasonably high. By contrast, when sequencing depth is low (Sim3, Sim5, and Sim6), the baseline ARI is substantially lower, consistent with earlier observations. In these scenarios, achieving a reasonable level of sensitivity or specificity is necessary for ssJSD methods to outperform the baseline (Figure 4, bottom panel, for Sim2, and Supplementary Figures S24-S28, bottom panels, for the rest of the scenarios).

### 2.5 ssJSD separates five known human cell types

We considered a dataset consisting of five human cell lines (H1Esc, HAP1, GM12878, HFF, and IMR90), available through the 4D Nucleome data portal (https://data.4dnucleome.org/). This dataset has served as a widely used benchmark for scHiC clustering, with methods such as topic modeling [29], BandNorm and scVI-3D [30], Higashi [20], and HiCENT [31] applied to the full dataset of thousands of cells. Due to the computational cost of HiCImpute, we selected 100 cells from each type with the highest sequencing coverage but excluding two H1Esc cells with exceptionally high read counts (over 1 million).

The datasets were binned at 1 Mb resolution. HiCImpute [21] was applied to identify SZs, which were then used to construct the sparsity matrix for each cell. Briefly, we computed the probability that each LP is an SZ. Then, the 95th percentile of the collection of these probabilities for the LPs with positive observed counts was used as the threshold to guard against false positives [23]. That is, an observed zero was classified as an SZ if its probability exceeded the threshold; otherwise, it was considered a DO.

When cells were ordered by sequencing depth within each of the five cell types, we observed that the proportion of OZs was strongly dependent on sequencing depth, with a larger proportion of OZs in cells with lower sequencing coverage (Supplementary Figure S29(a)). This observation underscores the widely held notion that many OZs are dropouts, which are believed to have arisen from insufficient sequencing depth. Indeed, the dropout proportion increases as sequencing depth decreases, mirroring the patterns of OZs. In contrast, the proportion of SZs appears to be largely invariant to sequencing depth, consistent with the notion that their origin is owing to their intrinsic genomic structure and biological mechanisms [22]. These findings provide additional support for the validity of the HiCImpute results. Interestingly, IMR90 exhibited a much higher proportion of structural zeros among all observed zeros compared to the other four cell types.

We considered four versions of the data when applying ssJSD: V1 — the sparsity matrix was constructed by treating all OZs as SZs; V2 — the sparsity matrix was constructed using SZs inferred from HiCImpute; V3 — in addition to constructing the SZ-induced sparsity matrix, the contact frequency matrix was improved by replacing DOs with HiCImpute values; and V4 — similar to V3, but with positive counts also replaced by HiCImpute values.

For comparison, we also applied JSD to three versions of the data: the observed, GK-improved, and RW-improved contact frequency matrices. Because of the sparsity of singlecell data and the fact that only spatial structural information was used, applying JSD to these three data versions did not yield clear separation of the five known cell types, whether based on the dissimilarity matrix or the 2D t-SNE projection (Figure 5a). These results echo those that we saw with the simulated data in the previous two subsections. Applying two ssJSD variants with V1 (treating all OZs as SZs) led to only minor improvement, which is unsurprising since a significant proportion (about half in four of the five cell types) of the OZs are DOs rather than SZs. In contrast, applying ssJSD to the other three data versions (V2-V4) that incorporated the HiCImpute-derived SZ-induced sparsity matrix resulted in marked improvements in separating the five cell types (Figure 5b for V3 and Supplementary Figure S30 for V2 and V4). Although a direct comparison with existing methods [29, 30, 20, 31] is not straightforward given the differences in the numbers of cells used, ssJSD with HiCImpute-derived SZ annotations achieves near-perfect separation of the five cell types judging from the four assessment criteria results (Supplementary Figure S31(a)).

**Figure 5:**
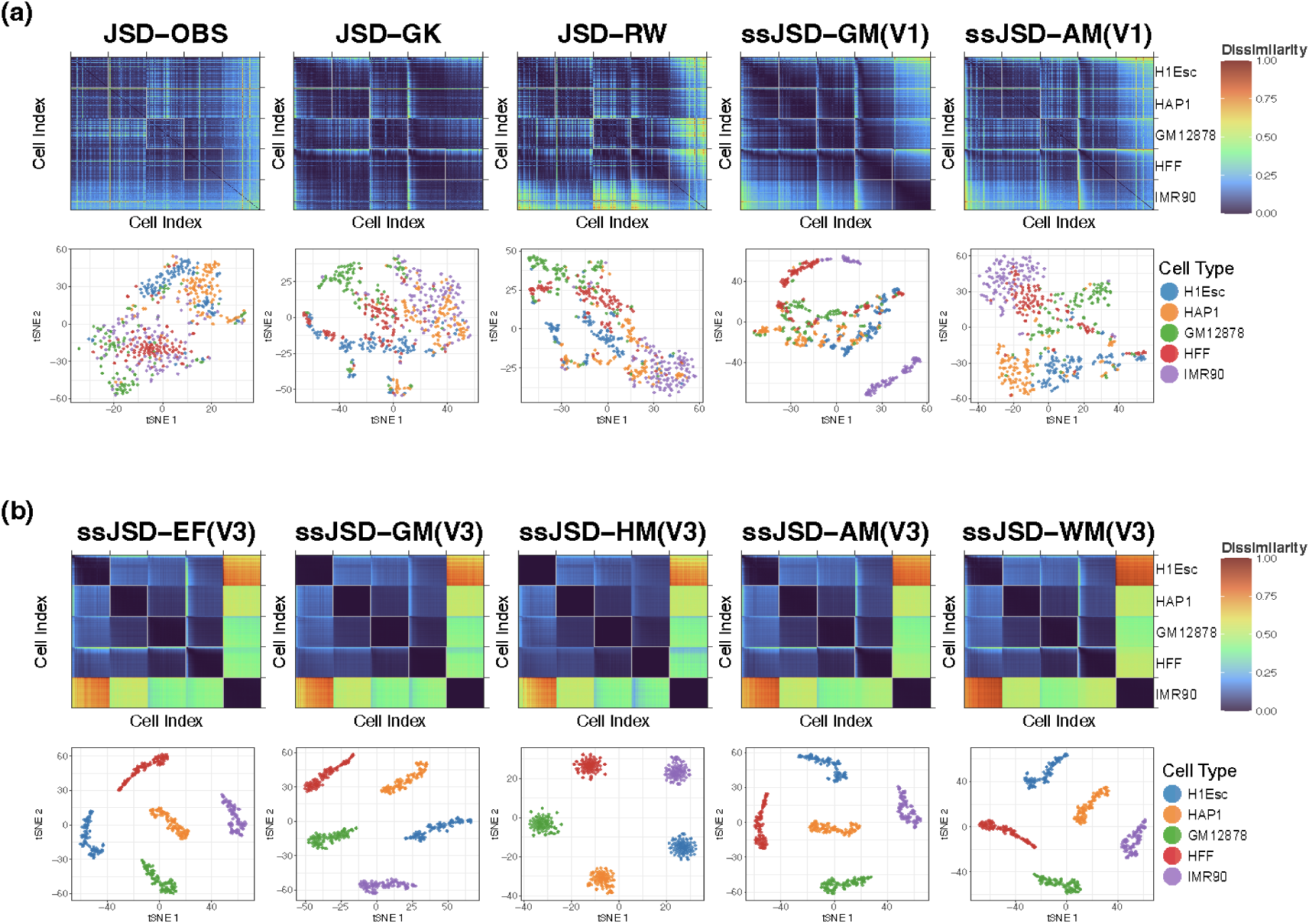
Results from analyzing the five human cell types data based on heatmap and 2D tSNE visualization. (a) Five methods without utilizing inferred SZ information; (b) Five data-fusion methods utilizing inferred SZ annotation and imputed values for DOs based on HiCImpute.

### 2.6 ssJSD delineates subtypes of neurons in the prefrontal cortex

We analyzed a second real dataset containing 4238 cells from 14 cell types in the prefrontal cortex [32] (accession number: GSE130711), collected from two donors (aged 21 and 29 years) at different time points, resulting in five batches. These subtypes are located in different cortical layers and are therefore expected to be separable. Inference from HiCImpute revealed a pattern similar to that observed in the previous dataset: the proportion of DOs increases as sequencing depth decreases, mirroring the trend for OZs. In contrast, the proportion of SZs remained stable across sequencing depths, further supporting the validity of HiCImpute’s results (Supplementary Figure S29(b)). Interestingly, although the proportion of OZs are similar to those in the five-human-cell-lines dataset, the proportions of SZs are much lower.

Applying JSD to four versions of the data without utilizing imputed SZ information—observed, RW- and GK-improved, and treating all OZs as SZs—yielded broadly similar results. Two large blocks are discernible (Figure 6a), one corresponding to neuronal cells (excitatory neurons—L2/3, L4, L5, L6 and inhibitory neurons—Pvalb, Sst, Vip, Ndnf) and the other to non-neuronal cells (Astro, OPC, ODC, MG, Endo, MP). However, the individual cell types within each block remained indistinguishable, reflecting the limitations of ignoring SZ information entirely (JSD methods) or naively treating all observed zeros as SZs (ssJSD-GM with V1 data). In contrast, ssJSD-GM applied to V2 data successfully separated all 14 cell types (Figure 6b). Moreover, incorporating additional imputed data from HiCImpute increased the between-subtype contrast, making the block structure clearer, especially when comparing V3 to V2. The zoomed panel of excitatory neurons from ssJSD-GM (V3) further reveals batch-induced substructure (Figure 6b, top-right plot). Specifically, cells in batch 181218 21yr are much more similar among themselves within each subtype compared to cells from the other four batches. Despite the batch effect, the between-subtype dissimilarity among L2/3, L4, L5, and L6 remains substantially larger than the within-subtype variation when all cells are considered together. For all four late-fusion variants, the three SZ-informed data versions (V2-V4) achieved similarly favorable criterion values, which are much better than those acieved by the early fusion approach or those without using SZ information (Supplementary Figure S31(b)). Further, the heatmaps and the 2D projection results are largely robust across all ssJSD methods (Supplementary Figure S32).

**Figure 6:**
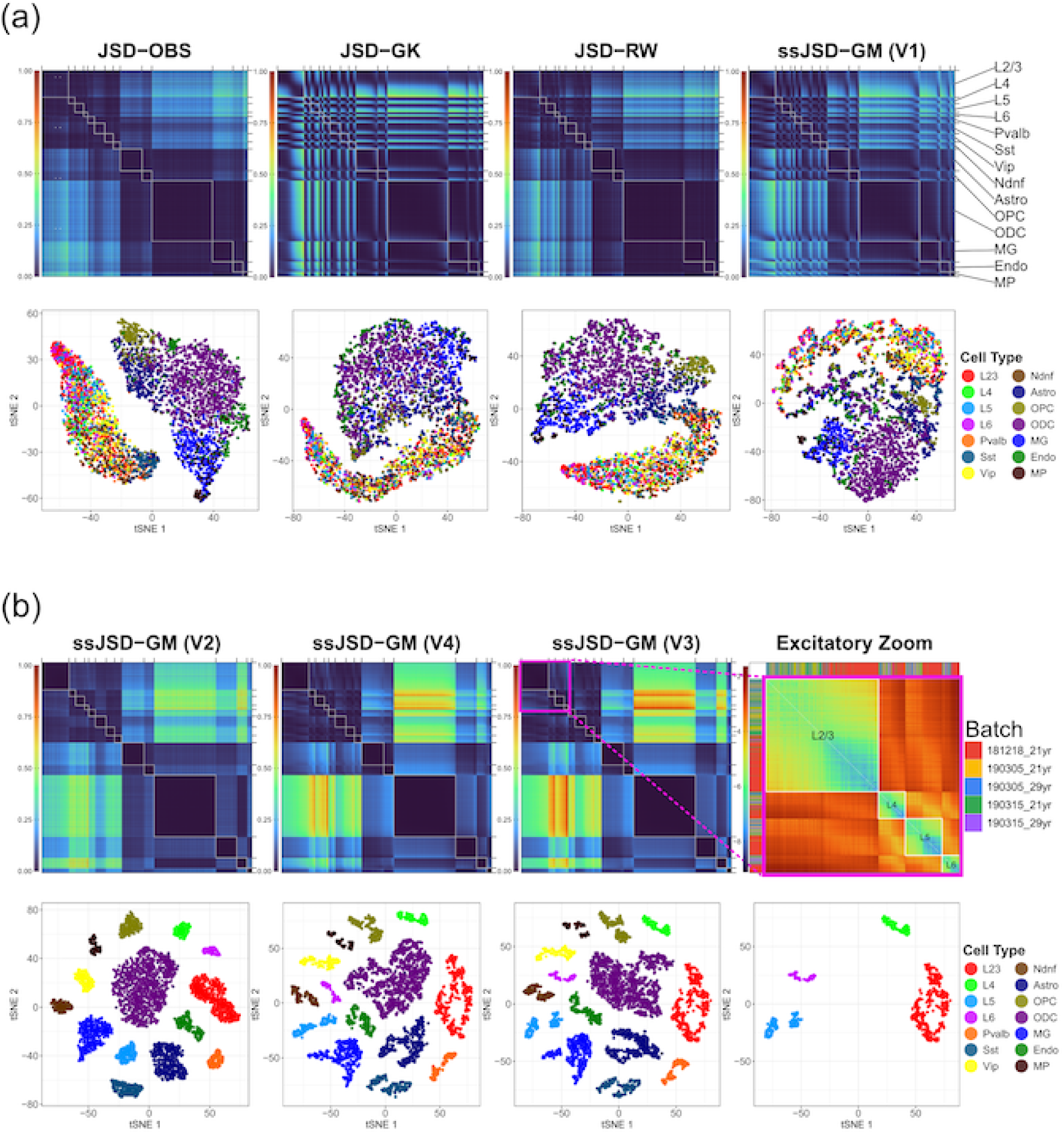
Results from analyzing the human prefrontal cortex data. In each panel, the top row shows the dissimilarity heatmaps and the bottom row shows the corresponding 2D tSNE visualizations. (a) Four methods without utilizing inferred SZ information; (b) ssJSD-GM applied to three versions of the data utilizing varying levels of results from HiCImpute—V2, V4, V3. The right most plot shows an enlarged heatmap of the excitatory neuron submatrix (L2/3, L4, L5, L6) from ssJSD-GM (V3) with color representing log(Dissimilarity); color bars on the left and top of the zoomed panel indicate the batch identity of each cell.

Existing methods applied to this dataset using scHiC contact matrices alone struggle to fully resolve all 14 cell types. BandNorm and scVI-3D [30] and Higashi [20] can broadly separate excitatory neurons, inhibitory subtypes, and non-neuronal cell types, but fail to distinguish individual excitatory neuronal layers (L2/3, L4, L5, and L6) from one another. The only SZ-informed method, szKendall [23], also achieves near-perfect separation of all 14 cell types; however, its *O*(*K*^2^*L*^2^) complexity renders it computationally prohibitive for scaling up, where *L* is the number of LPs and *K* the number of cells. In contrast, ssJSD achieves the same level of separation with *O*(*KL*+*K*^2^*B*) overall complexity through bandwise aggregation into *B ≪ L* genomic distance bands, substantially reducing the computational cost (Supplementary Figure S33).

## 3 Discussion

In this study, we introduced ssJSD, a novel dissimilarity framework that incorporates both spatial and sparsity information for single-cell HiC analysis. Specifically, ssJSD explicitly incorporates structural zero information alongside contact frequency profiles. By distinguishing true biological absences of chromatin contacts from technical dropouts, ssJSD provides a biologically informed quantification of cell-to-cell dissimilarity.

Through extensive simulation studies, we benchmarked ssJSD against existing approaches, including JSD applied to raw or smoothed contact matrices. When true SZ annotations were available, ssJSD greatly outperformed baseline measures, achieving more accurate subtype separation even under high sparsity. Importantly, the advantage of ssJSD persisted after dimensionality reduction with UMAP and t-SNE, which not only preserved but often enhanced subtype separation. Furthermore, robustness analyses under varying levels of SZ/DO misclassification demonstrated that ssJSD remained stable with a reasonably good performing SZ-inference tool: even with as low as 70% sensitivity and 80% specificity, the ssJSD variants generally surpassed JSD, which ignores sparsity information. Our results also show that simply treating all observed zeros as SZs is not a satisfactory alternative, as improvement over the baseline is minimal, if any, since the proportion of dropouts among the observed zeros is typically substantial.

Comparing the early fusion strategy against the late fusion approaches, the results favor the latter, perhaps due to the independent contributions to dissimilarity measures from both spatial count patterns and structural zero profiles. Among the four late-fusion variants proposed, the performances are comparable for the real data analysis examples. The weighted-mean method (ssJSD-WM) appears to be the best overall performer under the idealized setting where all the SZs and DOs are correctly identified; therefore, it is a strong contender to pair with a highly reliable algorithm for delineating the observed zeros.

In real data applications, we analyzed two representative datasets. First, in a collection of five human cell lines, ssJSD clearly separated the major cell types, achieving near-perfect clustering when HiCImpute-inferred SZ annotations were incorporated. In contrast, JSD applied to raw or smoothed data failed to identify these groups. Second, in the human prefrontal cortex dataset, ssJSD successfully separated all 14 cell types including the individual excitatory neuronal layers (L2/3, L4, L5, L6), a result often unachievable without explicitly incorporating structural zero information. These results emphasize that SZ information contributes critically to uncovering biologically meaningful structure that would otherwise remain hidden when zeros are treated uniformly. Furthermore, it appears that, in addition to SZ annotation, replacing dropouts with their imputed values may further improve clustering accuracy.

While our findings establish the utility of ssJSD, several limitations remain. The effectiveness of our method depends on accurate SZ inference. In this study, we relied on the probability outputs from HiCImpute to delineate OZs into SZs and DOs. Alternatively, instead of finding an appropriate threshold for the delineation, one may use the probabilities directly to construct the sparsity matrices. Additionally, more advanced algorithms may further improve performance, and computational speedup is necessary to avoid the bottleneck. Second, our results indicate that late fusion generally yields superior performance when true SZ annotations are available, although its accuracy declines in the presence of SZ misspecification. In contrast, early fusion provides only modest improvements under certain parameter settings, yet delivers more stable results overall. Future work may explore more adaptive fusion schemes that dynamically weight SZ and contact-frequency contributions depending on data quality. Finally, although ssJSD can be implemented in an efficient algorithm, its computational efficiency can be further improved by reducing the size of the *P_CF_*and *P_SZ_* vectors if information from bands far away from the diagonals contains more noise than signal, especially by setting the length of the vector to correspond to the size of the smallest, rather than the largest chromosome.

Taken together, our study demonstrates that incorporating SZ information into dissimilarity measures substantially improves the analysis of scHiC data. ssJSD enhances clustering accuracy and increases robustness to sparsity by utilizing biologically meaningful substructures and spatial patterns across both simulated and real datasets. We evaluated ssJSD primarily in clustering applications, but its dissimilarity matrices may also be valuable for other downstream analyses, such as network-based embedding or integrative multimodal studies.

## 4 Methods

### 4.1 Building band-wise probability vectors

To quantify the chromatin interaction patterns, we denote 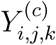 as the frequency of observed contact between loci *i* and *j* in chromosome *c* (*i* = 1*, . . . , B_c_ −* 1 and *j* = *i* + 1*, . . . , B_c_*) in cell *k* (*k* = 1*, . . . , K*), where a locus, in this context, is an equal-sized genomic region and *B_c_*is the number of bins in chromosome *c*. Following Liu et al. [12], we define a vector representation, *P_CF_* , of the contact frequencies (CF) for each band *s* in cell *k*:

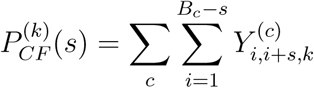

where *s* = 1, 2*, . . . , B* is the genomic-distance band, and *B* = max*_c_*(*B_c_*) *−* 1. Note that for chromosomes with *B_c_ < B*, 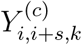 is not defined when *s ≥ B_c_*, and the contribution of such chromosomes to the sum is zero. Alternatively, one may set *B* = min*_c_*(*B_c_*) *−* 1 if bands far away from the diagonals are believed to contribute more noise than signal (also see Discussion). For each cell, this vectorized *P_CF_* (a probability vector after appropriate normalization) characterizes the aggregated contact pattern of the cell across all locus pairs (LPs) falling within diagonal band *s* parallel to the main diagonal of the contact matrix. Through aggregation, this probability vector of contact profile reduces variability while maintaining spatial information, since contact counts generally decrease as the distance between bins increases. In addition to observed contact frequencies, we assume the availability of cell-specific information that indicates whether each observed zero entry corresponds to a structural zero (SZ) that represents a true biological absence of chromatin interaction [21]. Specifically, let 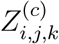 be a dichotomous indicator variable such that

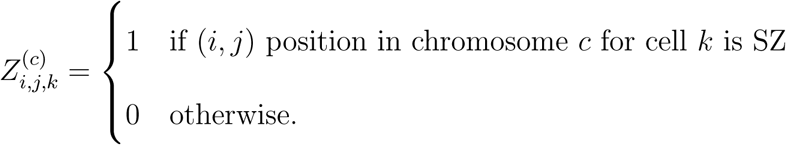

This binary labeling allows us to distinguish biologically meaningful zeros from technical dropouts. Following the same band-wise framework in constructing *P_CF_* , we define SZ-based feature vector 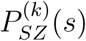 that summarizes the frequency of structural zeros in band *s* for cell *k*:

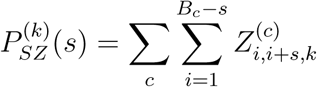

This reduced representation captures the spatial distribution of structural zeros across genomic distances, providing a complementary view of chromatin organization that focuses on biologically constrained absence of contact. Together with the *P_CF_* ’s, these SZ-based vectors *P_SZ_*’s enable the construction of more informed dissimilarity measures across single cells.

### 4.2 Combining the two vectors

We adopted two different strategies to combine the two vectors: early fusion and late fusion. In the early fusion approach, we first construct a concatenated vector, denoted by 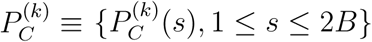, by normalizing each of 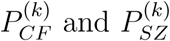 so that the sum of elements in each vector equals 1*/*2:

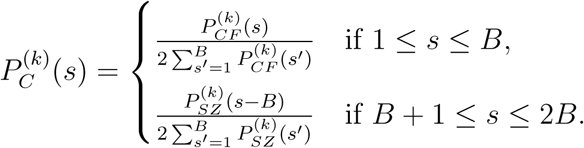

Then we compute JSD between the concatenated vectors of each pair of cells. Specifically, for any pair *k* and *k*^′^, we define

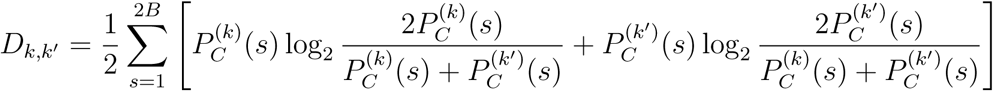

In the late fusion approach, each CF and SZ vector is normalized independently:

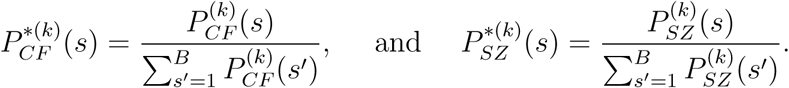

Then, we compute JSD separately based on each of the two normalized vectors. This results in two pairwise dissimilarity matrices 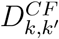 and 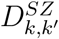.

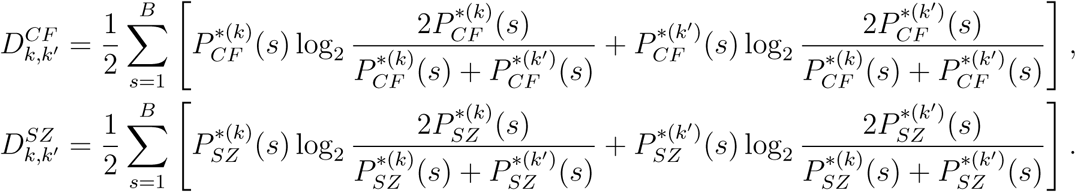

After computing the JSDs, we applied min-max normalization to each dissimilarity matrix to ensure that both are on the same scale before integration. Specifically, we define

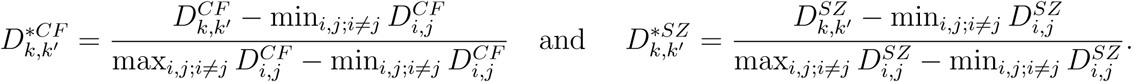

Finally, we merge 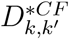 and 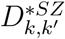 to integrate two complementary sources of information: one capturing dissimilarities in positive contact counts, and the other reflecting dissimilarities in structural zero patterns, both taking spatial information into consideration. We considered four ways of combining the two matrices to arrive at a joint one:

1. Geometricmean(GM

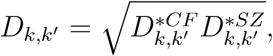

2. Harmonicmean(HM)

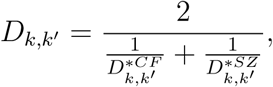

3. Arithmeticmean(AM)

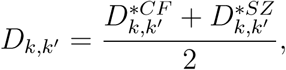

4.Weightedmean(WM)

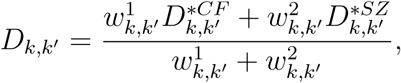

where 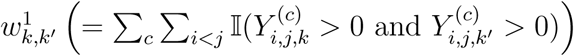 is the number of matched positive counts between cell *k* and *k*^′^, whereas 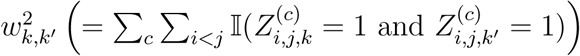 is the number of matched structural zeros between cell *k* and *k*^′^.

### 4.3 Simulation procedure for producing Sim1-6 data

The following procedure is adapted from the literature [14, 21] to produce the 12 sets of data corresponding to the six scenarios of parameter settings (Table 1) and the two designs. Specifically, for each subtype under each of the 12 settings, we generated single cell contact matrices for each subtype following a three-step procedure as follows.

**Step 1: Modeling Expected Contact Counts.** First, we modeled the expected contact frequency, *λ_ij_*, for every pair of loci (*i, j*) based on the 3D Euclidean distance (*d_ij_*) derived from the underlying structure. The log-expected count was defined as:

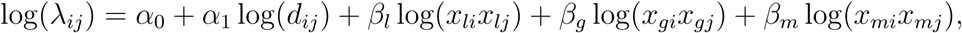

where *α*_0_ regulates sequencing depth and *α*_1_ controls the rate of distance decay. To introduce biological variability, we incorporated three covariates—representing fragment length (*x_l_*), GC content (*x_g_*), and mappability (*x_m_*)—sampled from uniform distributions: *x_li_ ∼* Uniform(0.2, 0.3), *x_gi_ ∼* Uniform(0.4, 0.5), and *x_mi_ ∼* Uniform(0.9, 1.0), with fixed coefficients *β_l_* = *β_g_* = *β_m_*= 0.9.

**Step 2: Identifying SZ Candidates.** Next, we established a pool of potential structural zeros. Pairs with low expected counts (*λ_ij_*) falling below the (*γ ×* 100)-th percentile were flagged as initial candidates. From this set, we randomly selected a subset of pairs with probability *τ*_1_ to serve as the candidate pool, denoted as *S_cand_*.

**Step 3: SZ Assignment.** Finally, to reflect cell-to-cell variability even within a subtype, the candidates in *S_cand_* were assigned based on two distinct categories:

- **Common SZ Assignment:** A fixed proportion *η* from the candidate pool *S_cand_* was designated as *Common SZs*. These pairs were universally assigned as structural zeros across all cells within the subtype.
- **Random SZ Assignment:** The remaining 1 *− η* proportion of *S_cand_* served as candidate for *Random SZs*. For each individual cell, every candidate in this set was independently assigned as a structural zero with a probability of *τ*_2_, resulting in approximately *τ*_2_ *×* 100% of these positions being set to zero for that specific cell.

Observed counts *Y_ij_*were then generated from a Poisson distribution with mean *λ_ij_*, setting *Y_ij_* = 0 if the pair was assigned as an SZ.

### 4.4 Four criteria for comparison of clustering performance

Following the clustering strategy where PAM is directly applied to the dissimilarity measure and Kmeans is used for projected dissimilarity measure, we used the same set of four metrics as in szKendall [23] to evaluate the performance of the results. Specifically, the Adjusted Rand Index (ARI) is used as an external metric to assess the accuracy of clustering results in recovering the ground truth cell types. For internal structural evaluation, we adopt the Within-cluster Sum of Squares percentage (WSS%), Average Silhouette Width (ASW), and the Minimum Isolation Score (MIS) to measure cluster cohesion and separation.

## Data Availability

The ssJSD R package, together with the prepared R data for the real and simulated data used in this study, are available on Github: https://github.com/osu-stat-gen/ssJSD.

## Acknowledgements

This research is supported in part by a grant from the National Institute of Health R01GM114142.

## Author Contributions

SL designed the study and supervised the project. SWL conducted the research. Both authors wrote the manuscript.

## Additional information

Correspondence and requests for materials should be addressed to SL.

