## Supplementary figures for "ssJSD: A Fusion of Sparsity and Spatial Information for HiC Single-Cell Clustering"

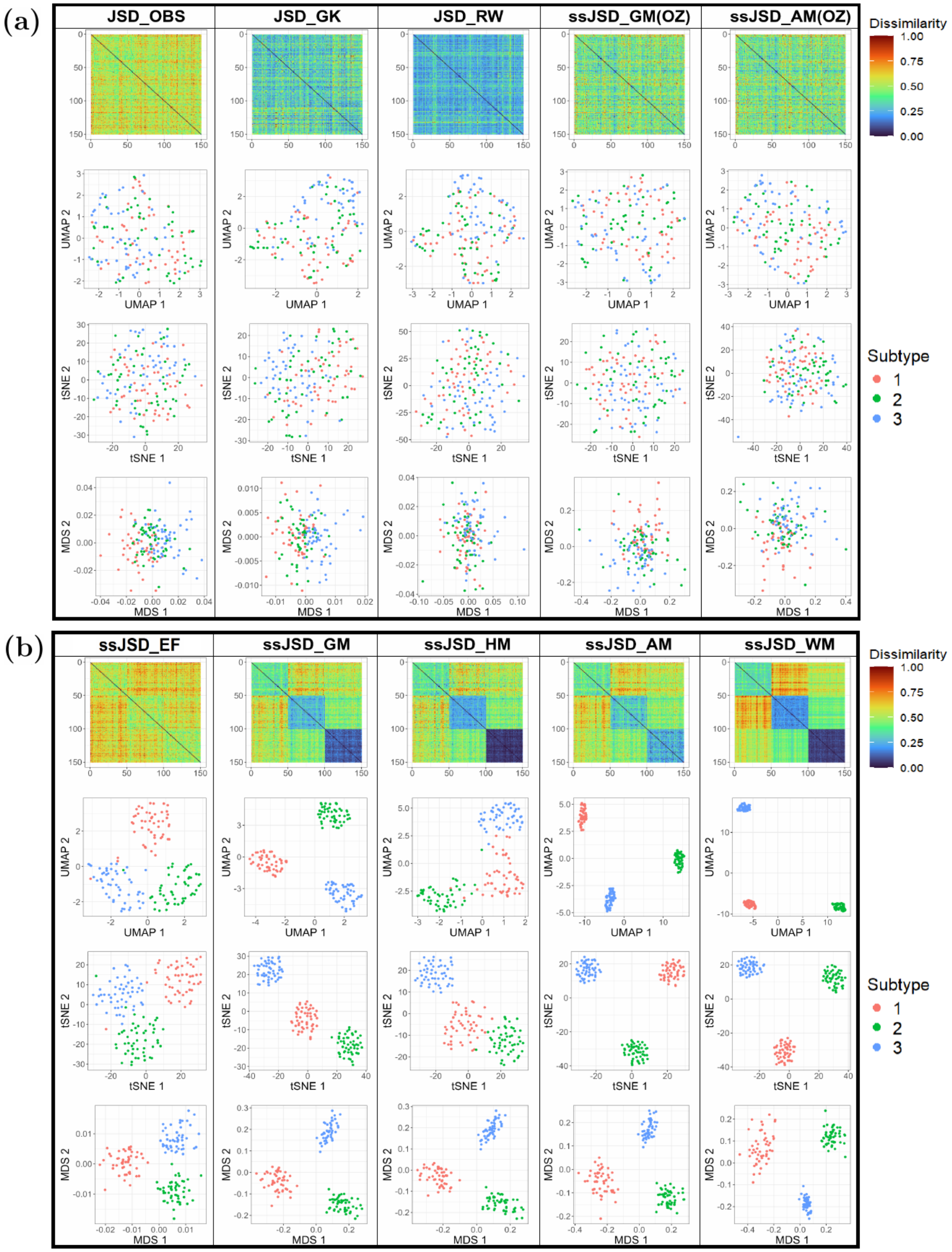


Figure S1: Comparison of ten methods in terms of heatmap and 2D projections of the dissimilarity matrix for Design 1 Sim1. (a) Five methods that do not use correct SZ information; (b) Five proposed methods that use SZ information.

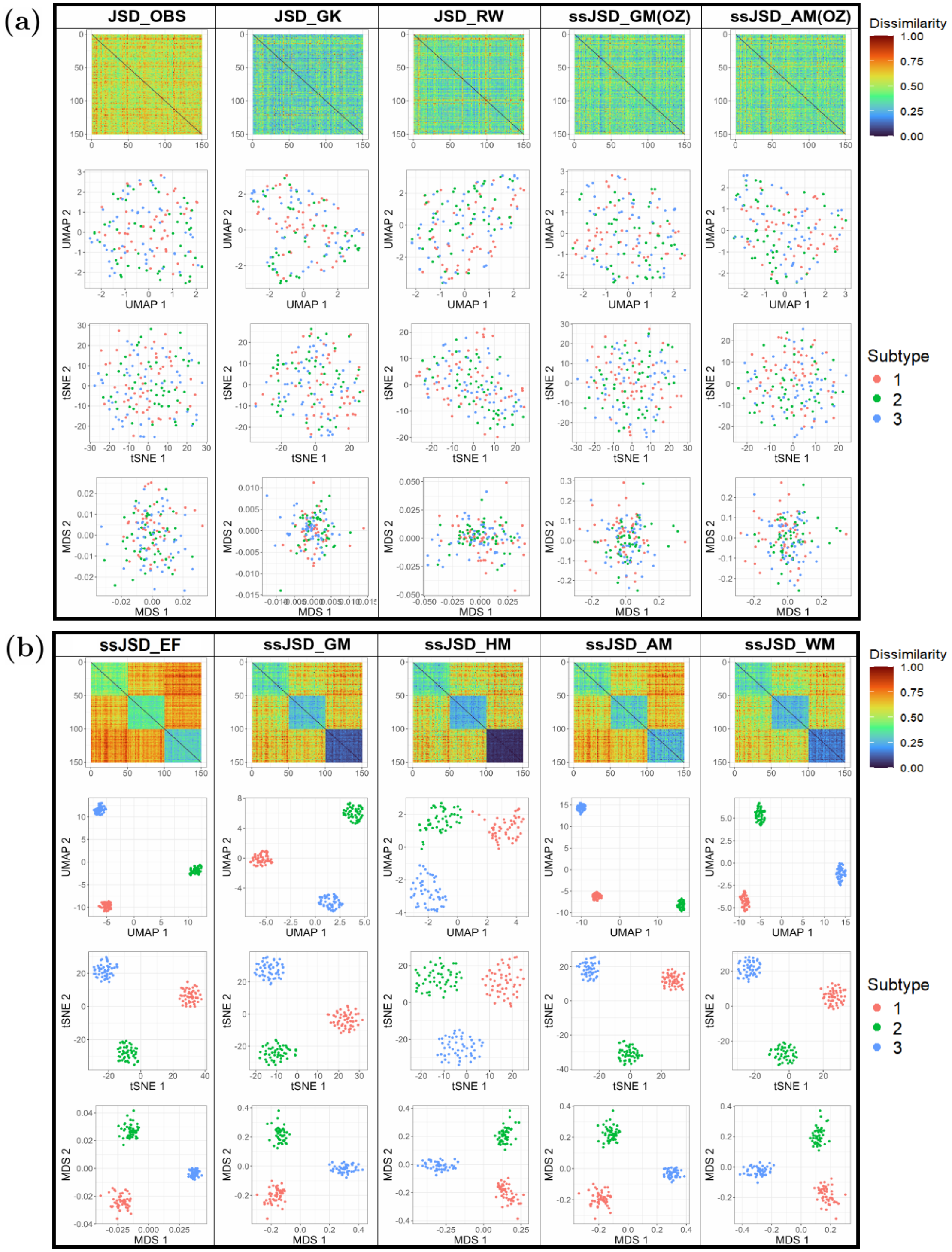


Figure S2: Comparison of ten methods in terms of heatmap and 2D projections of the dissimilarity matrix for Design 1 Sim2. (a) Five methods that do not use correct SZ information; (b) Five proposed methods that use SZ information.

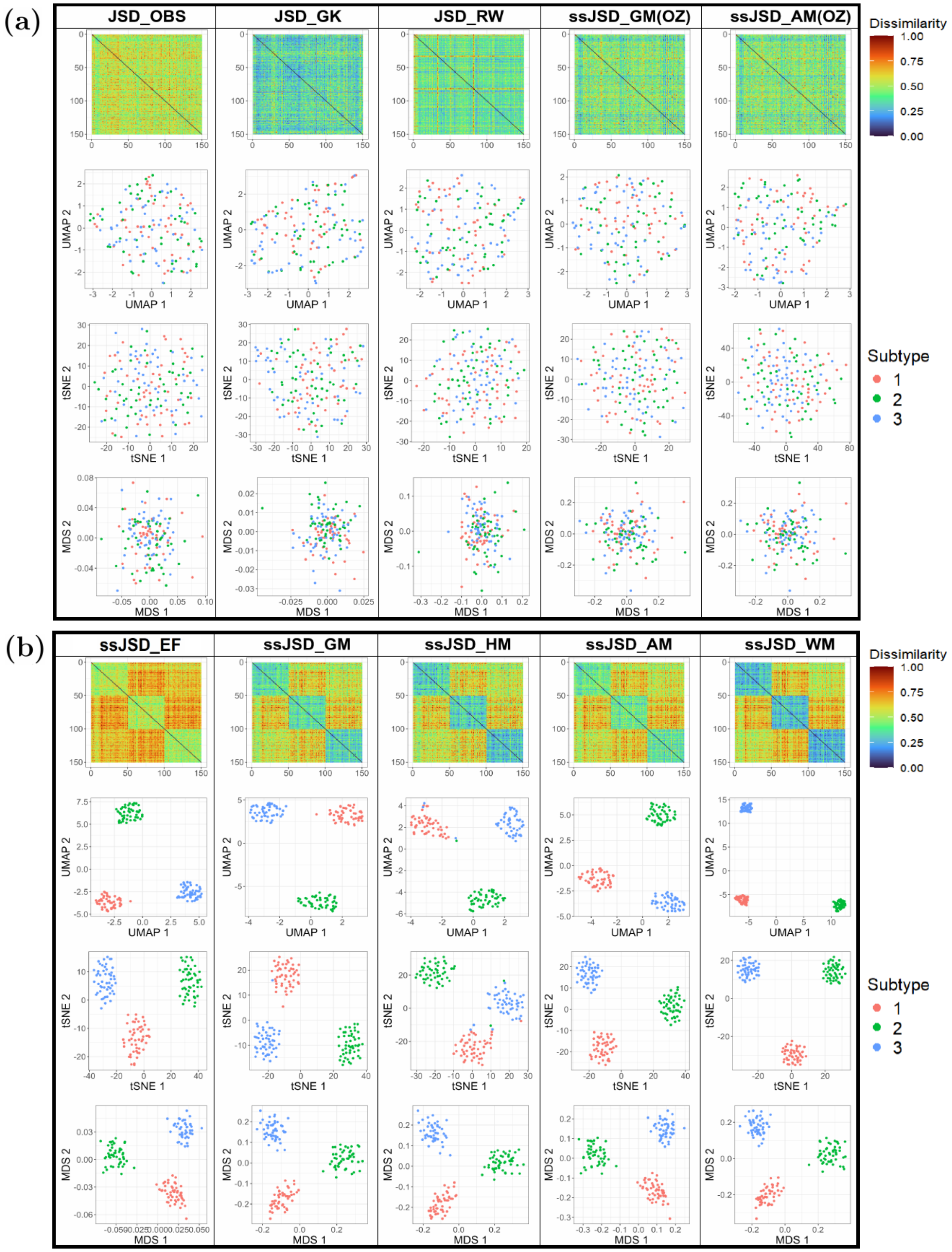


Figure S3: Comparison of ten methods in terms of heatmap and 2D projections of the dissimilarity matrix for Design 1 Sim3. (a) Five methods that do not use correct SZ information; (b) Five proposed methods that use SZ information.

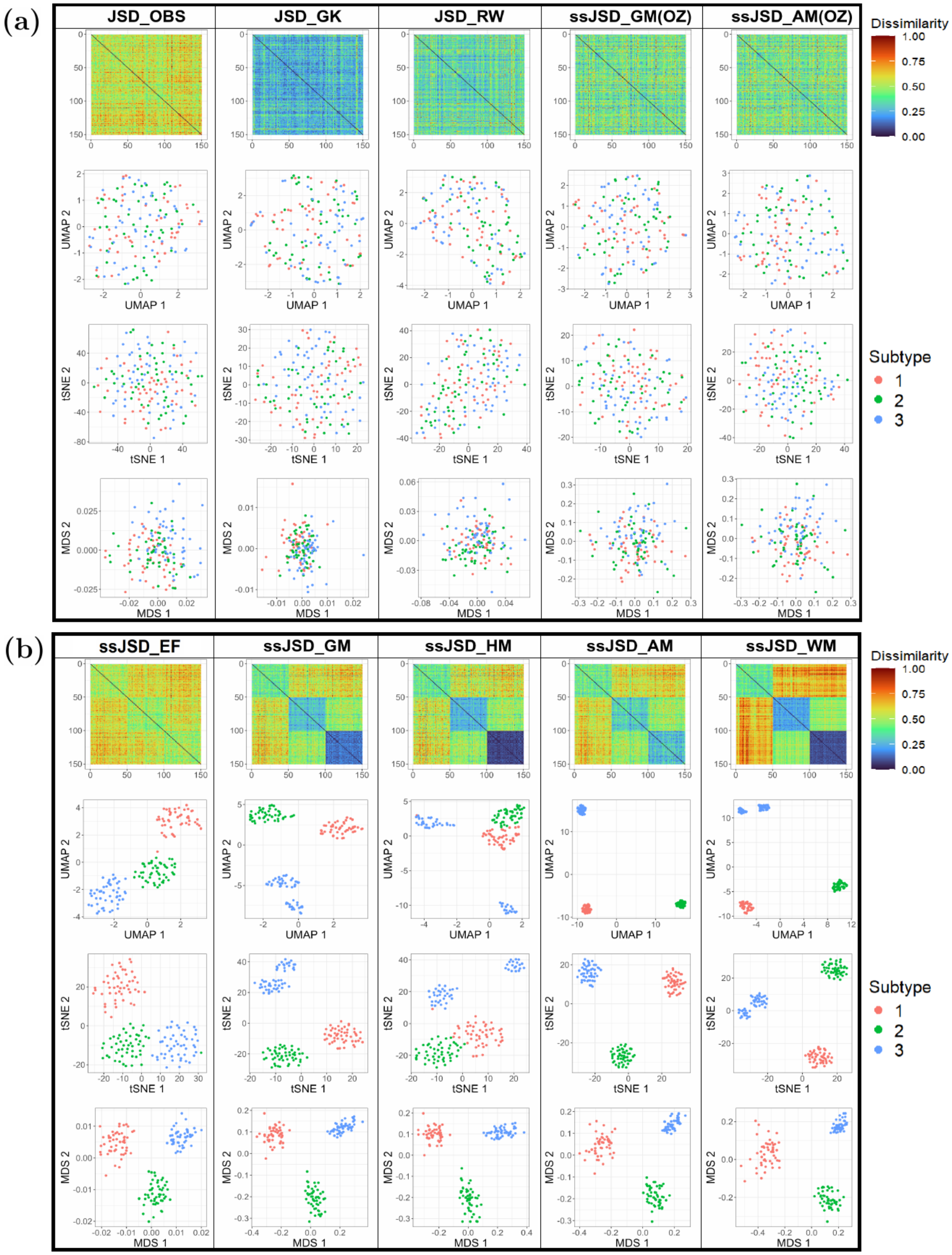


Figure S4: Comparison of ten methods in terms of heatmap and 2D projections of the dissimilarity matrix for Design 1 Sim4. (a) Five methods that do not use correct SZ information; (b) Five proposed methods that use SZ information.

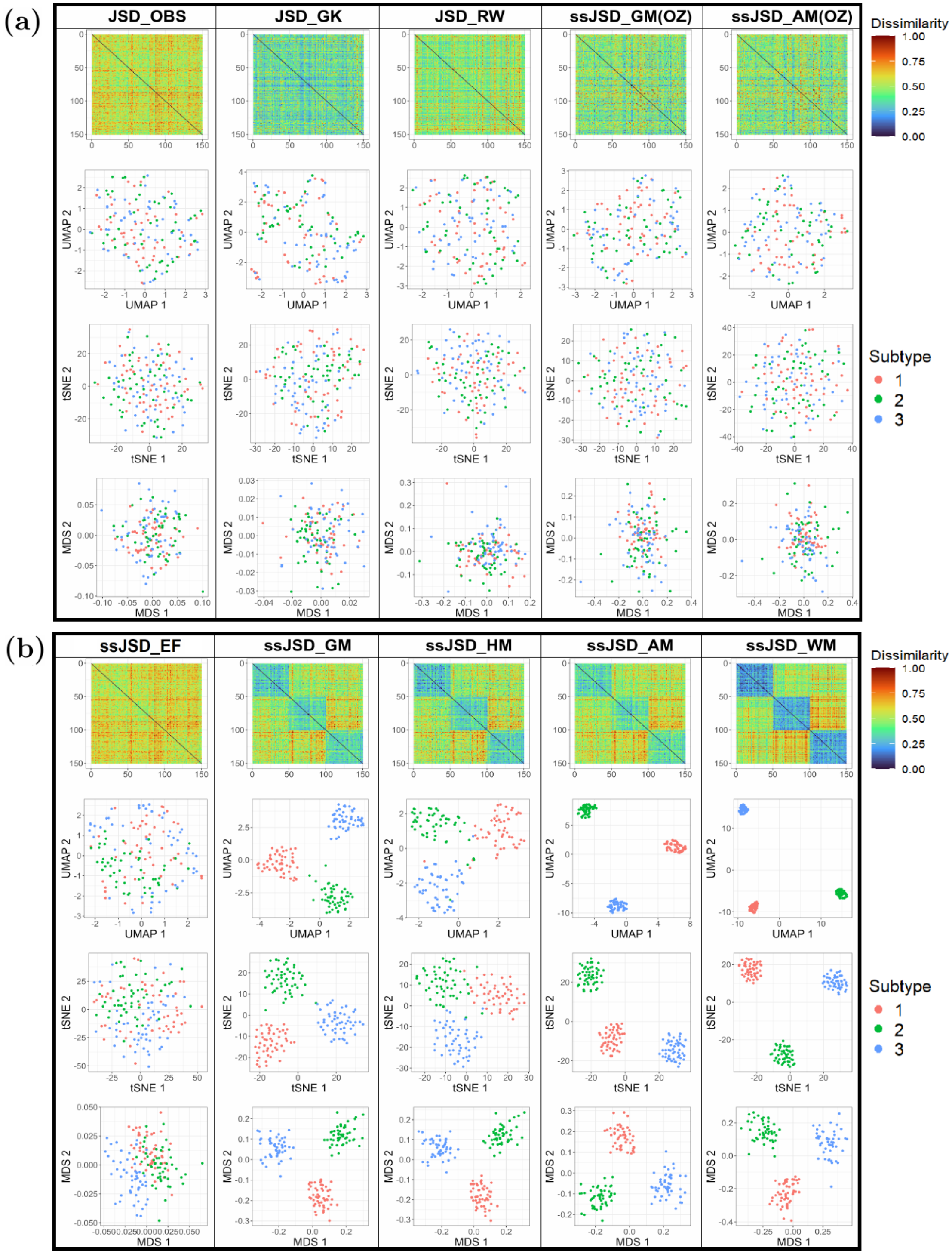


Figure S5: Comparison of ten methods in terms of heatmap and 2D projections of the dissimilarity matrix for Design 1 Sim5. (a) Five methods that do not use correct SZ information; (b) Five proposed methods that use SZ information.

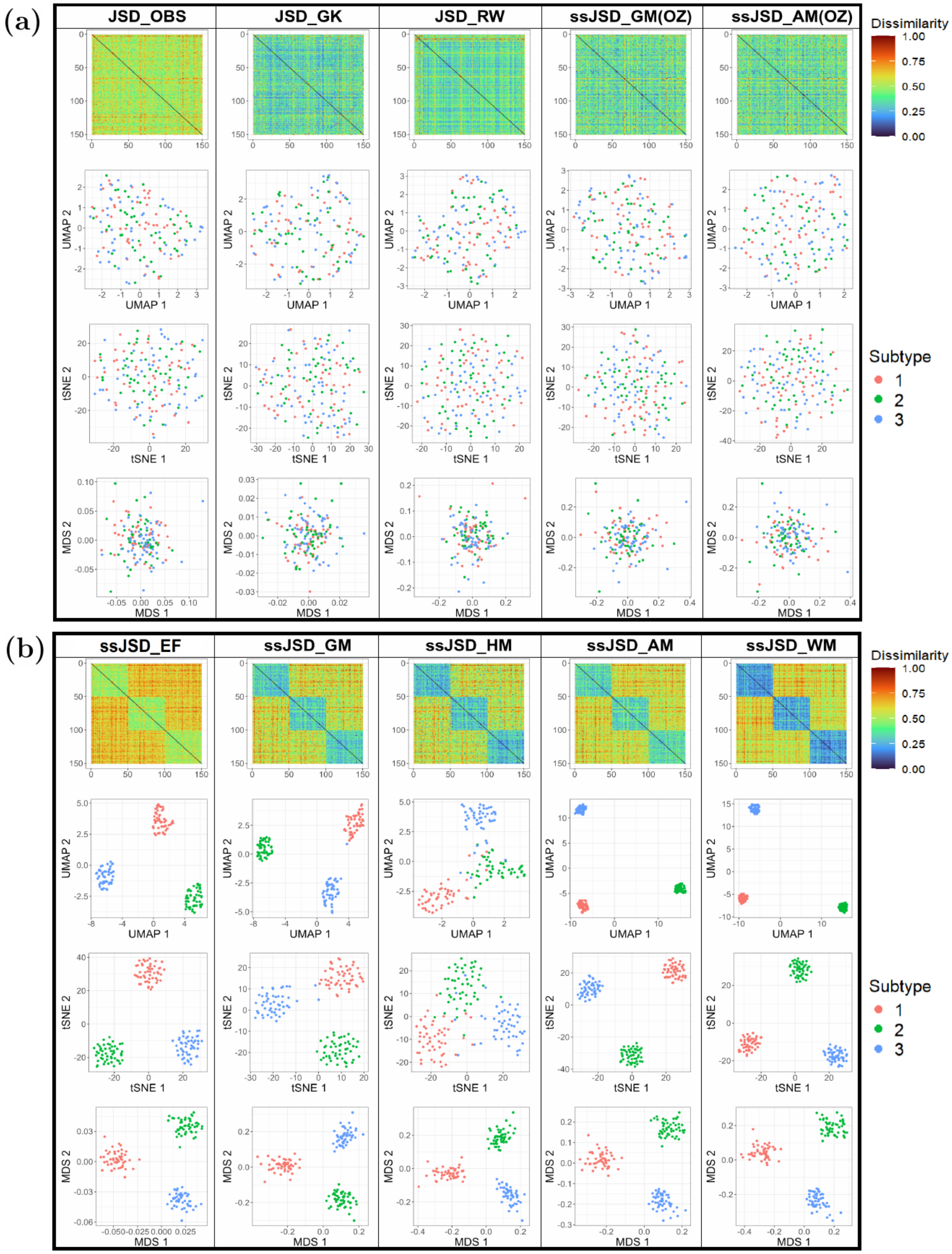


Figure S6: Comparison of ten methods in terms of heatmap and 2D projections of the dissimilarity matrix for Design 1 Sim6. (a) Five methods that do not use correct SZ information; (b) Five proposed methods that use SZ information.

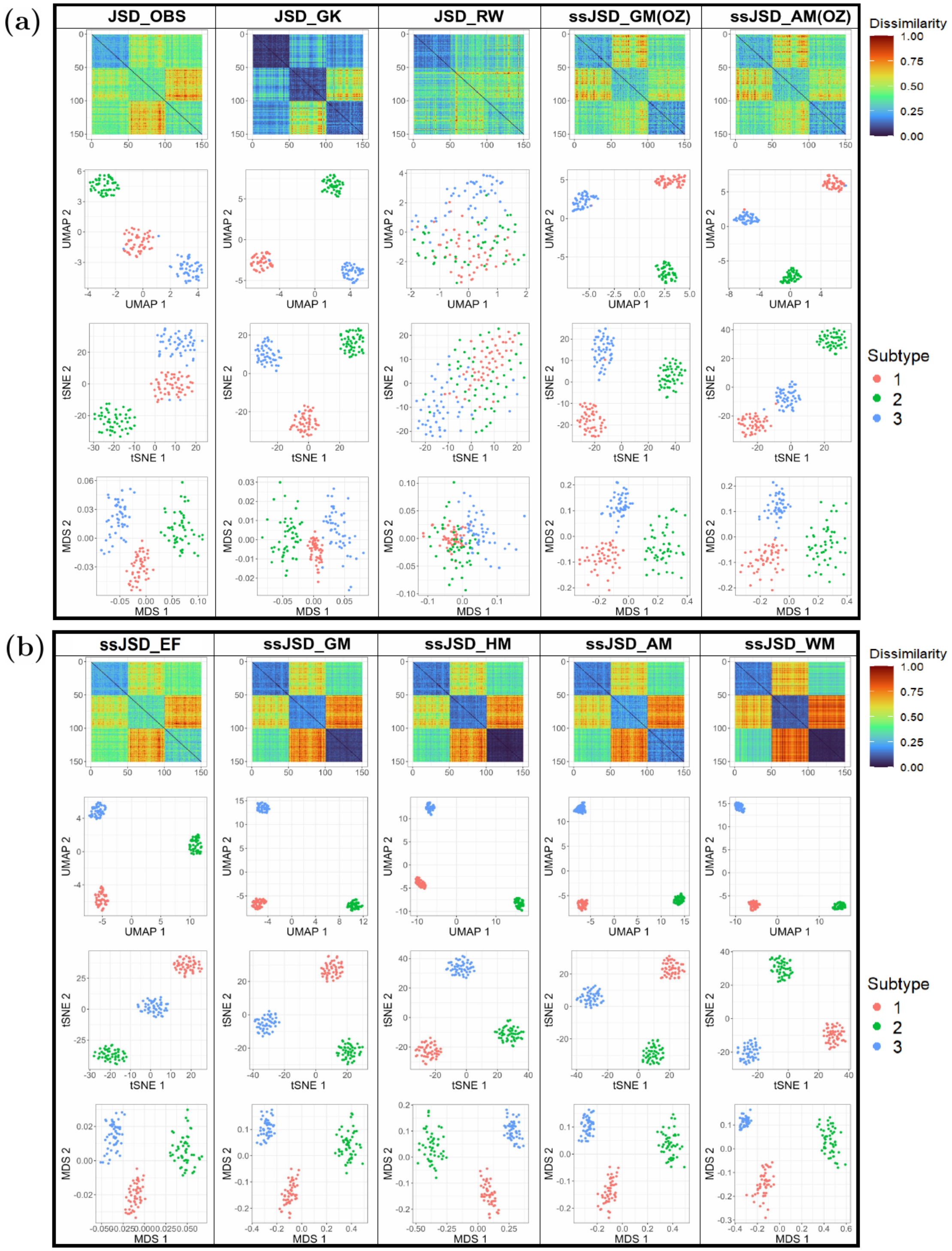


Figure S7: Comparison of ten methods in terms of heatmap and 2D projections of the dissimilarity matrix for Design 2 Sim1. (a) Five methods that do not use correct SZ information; (b) Five proposed methods that use SZ information.

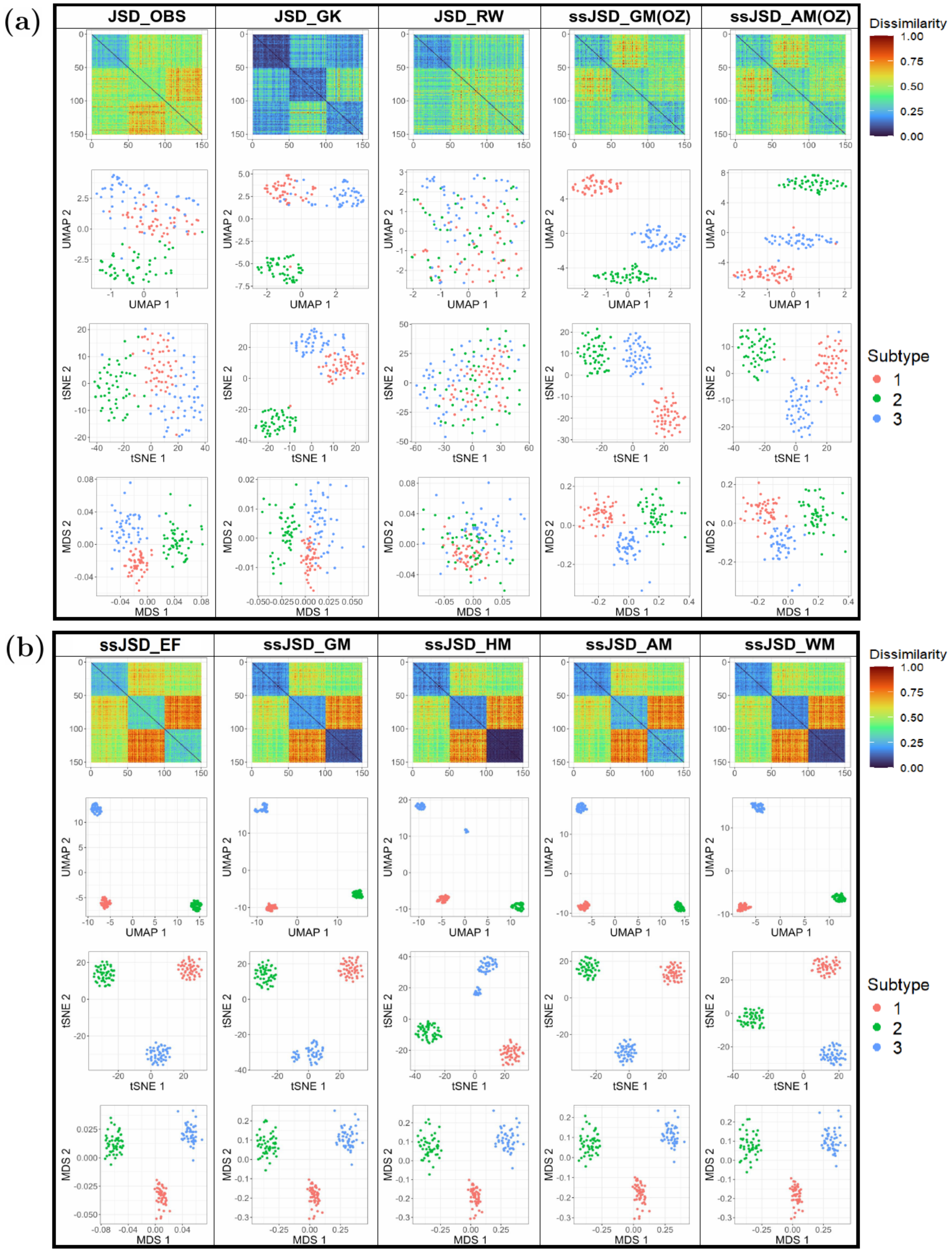


Figure S8: Comparison of ten methods in terms of heatmap and 2D projections of the dissimilarity matrix for Design 2 Sim2. (a) Five methods that do not use correct SZ information; (b) Five proposed methods that use SZ information.

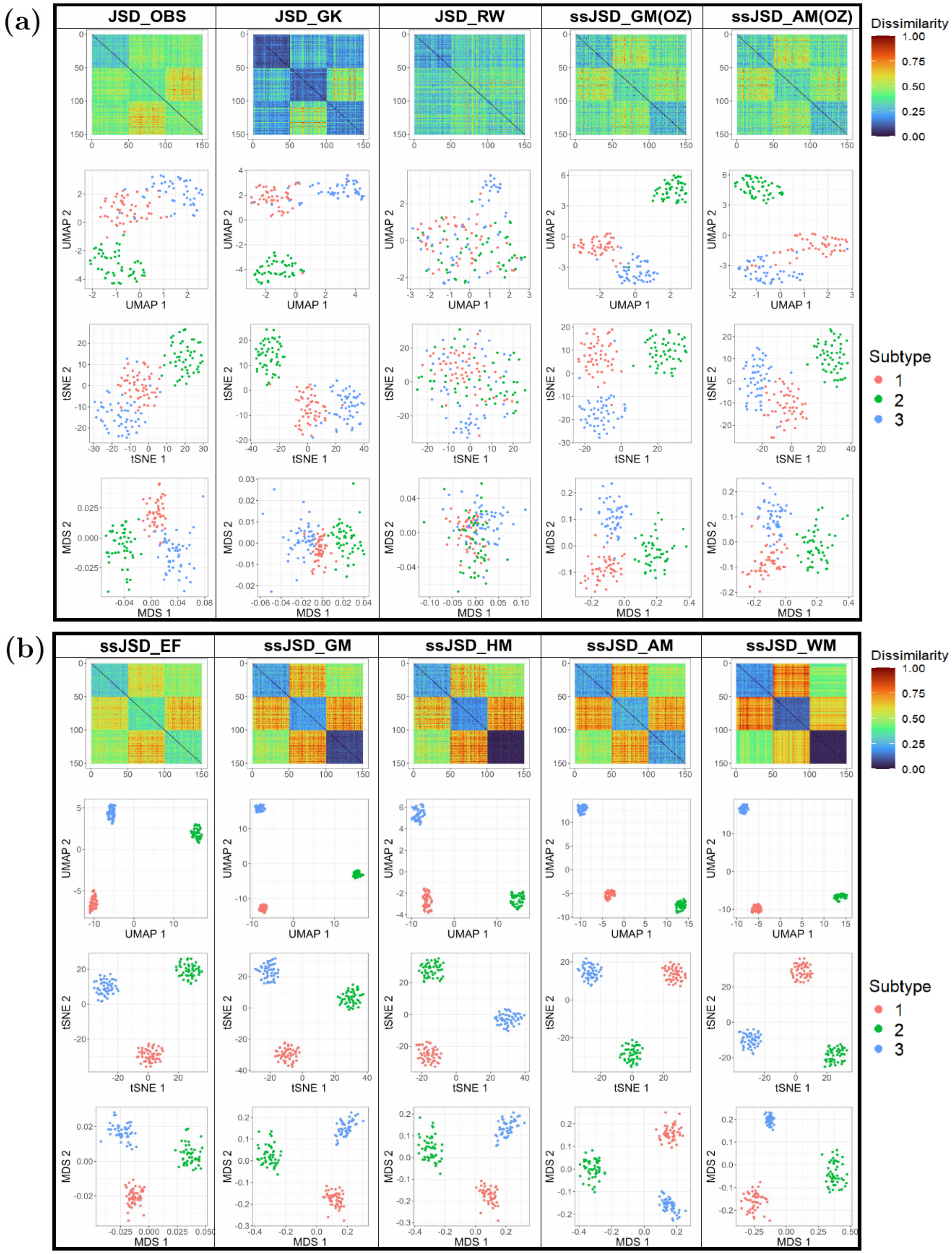


Figure S9: Comparison of ten methods in terms of heatmap and 2D projections of the dissimilarity matrix for Design 2 Sim4. (a) Five methods that do not use correct SZ information; (b) Five proposed methods that use SZ information.

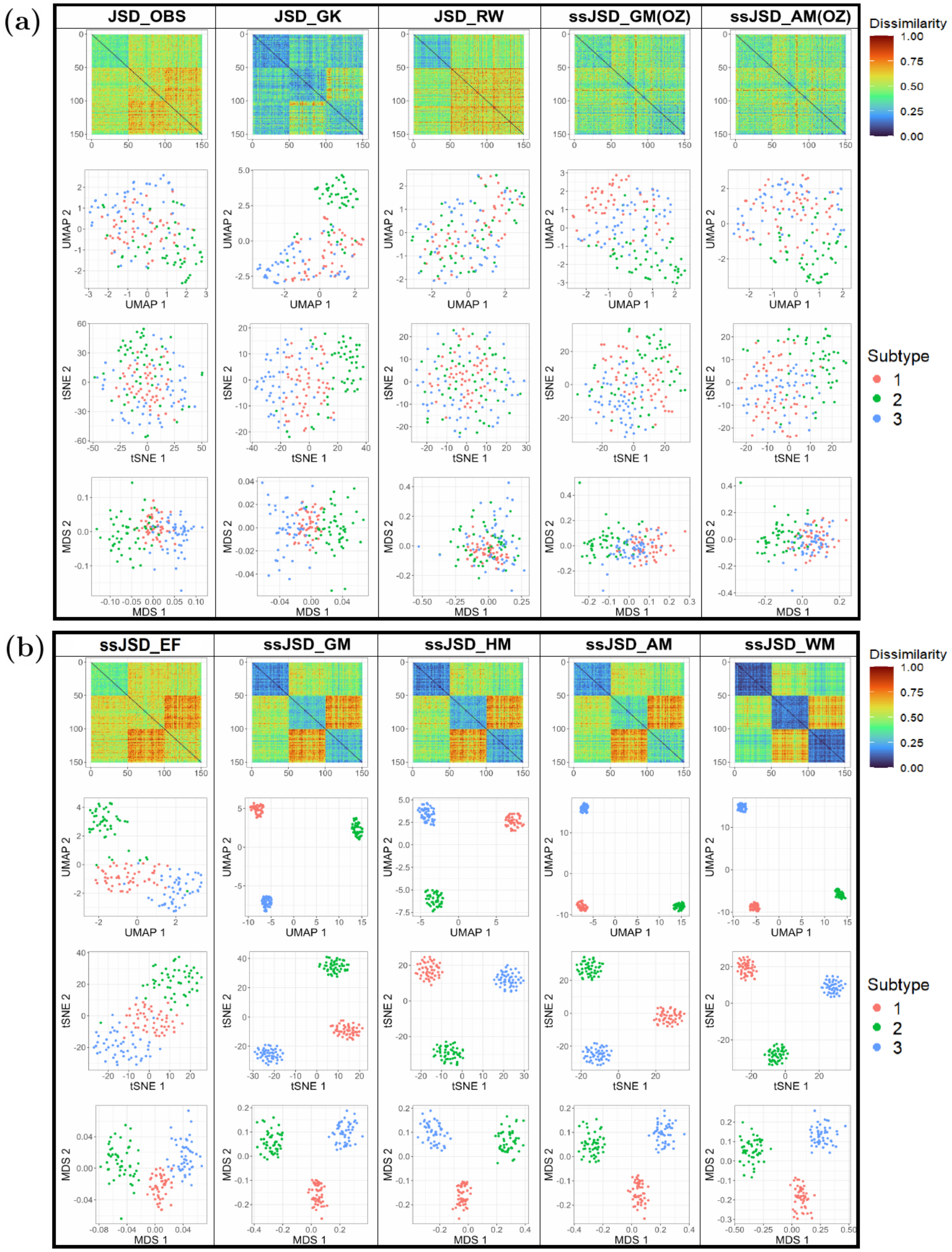


Figure S10: Comparison of ten methods in terms of heatmap and 2D projections of the dissimilarity matrix for Design 2 Sim5. (a) Five methods that do not use correct SZ information; (b) Five proposed methods that use SZ information.

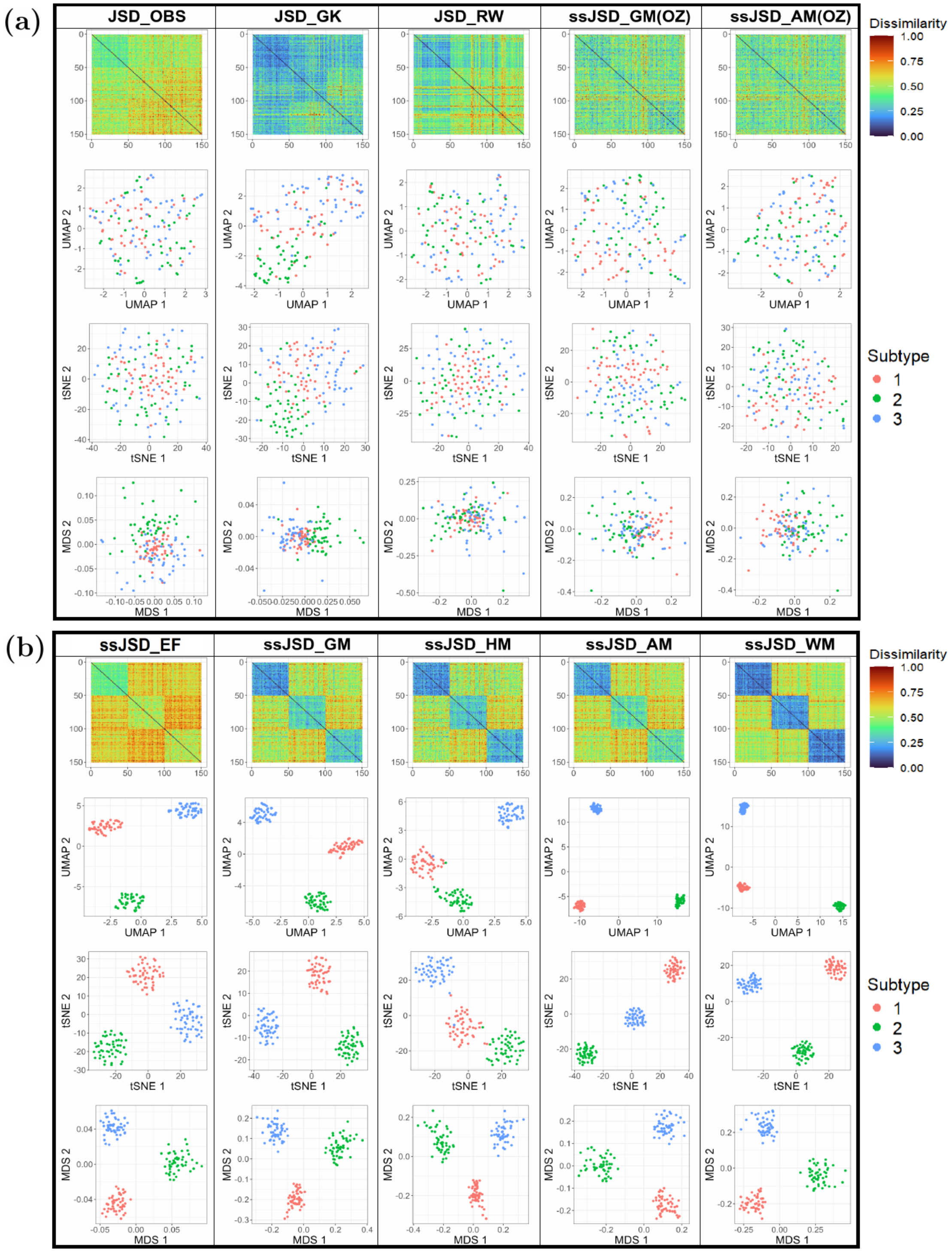


Figure S11: Comparison of ten methods in terms of heatmap and 2D projections of the dissimilarity matrix for Design 2 Sim6. (a) Five methods that do not use correct SZ information; (b) Five proposed methods that use SZ information.

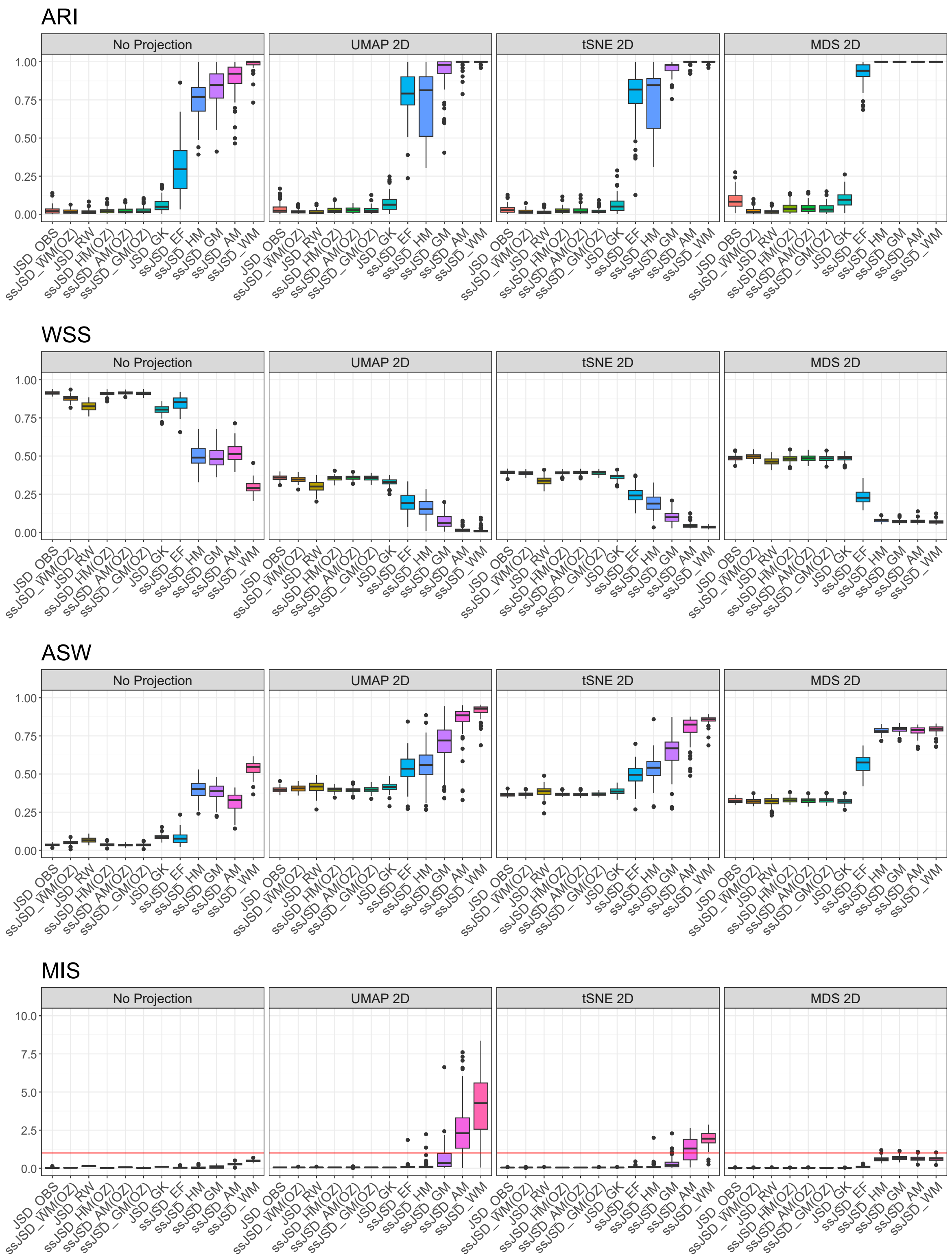


Figure S12: Comparison of the performance of 12 methods based on four clustering criteria—ARI, WSS, ASW, and MIS—across four data representations: the dissimilarity matrix (no projection), or three 2D projections (UMAP, tSNE, and MDS). The boxplots show the results across 100 datasets under Design 1 Sim1.

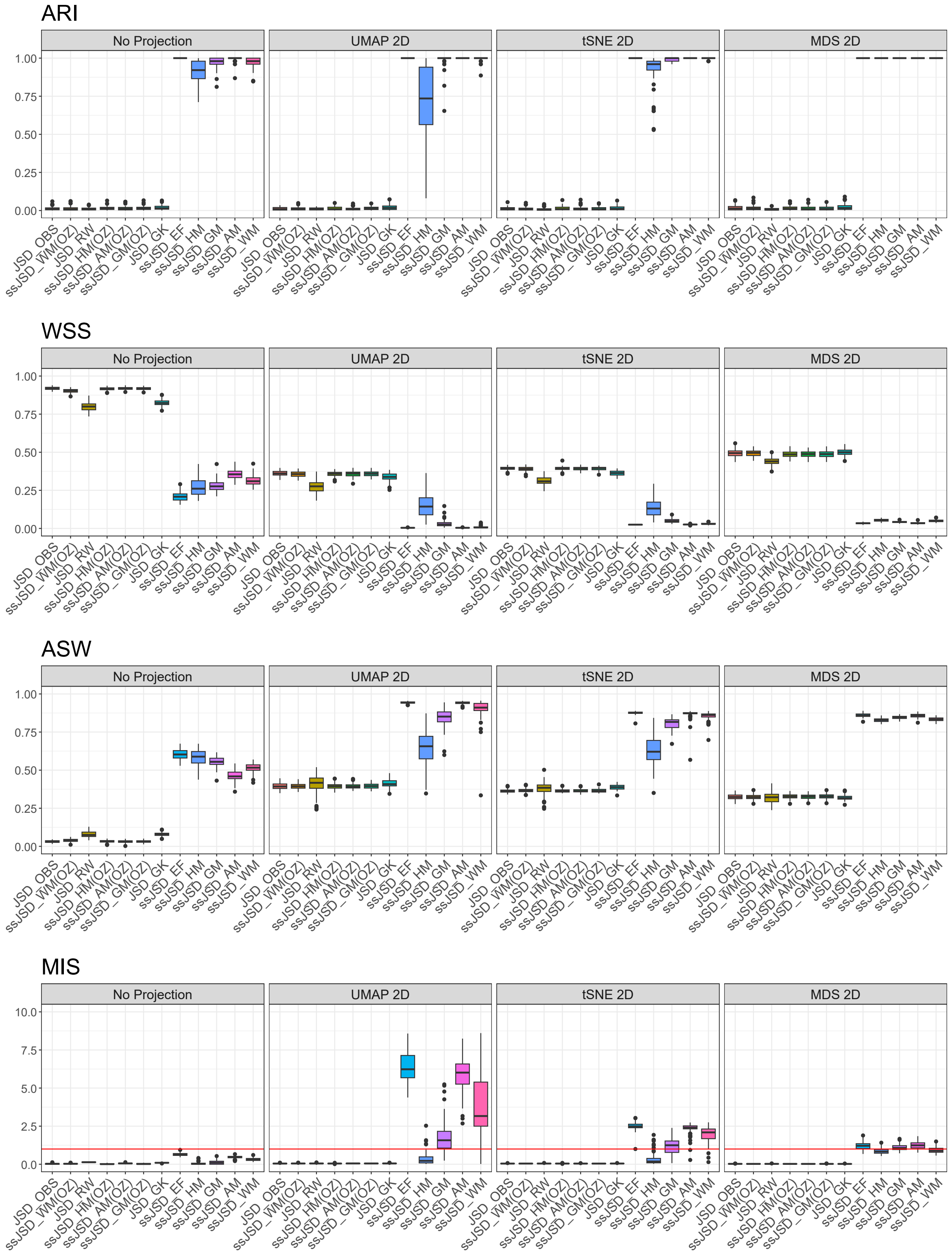


Figure S13: Comparison of the performance of 12 methods based on four clustering criteria—ARI, WSS, ASW, and MIS—across four data representations: the dissimilarity matrix (no projection), or three 2D projections (UMAP, tSNE, and MDS). The boxplots show the results across 100 datasets under Design 1 Sim2.

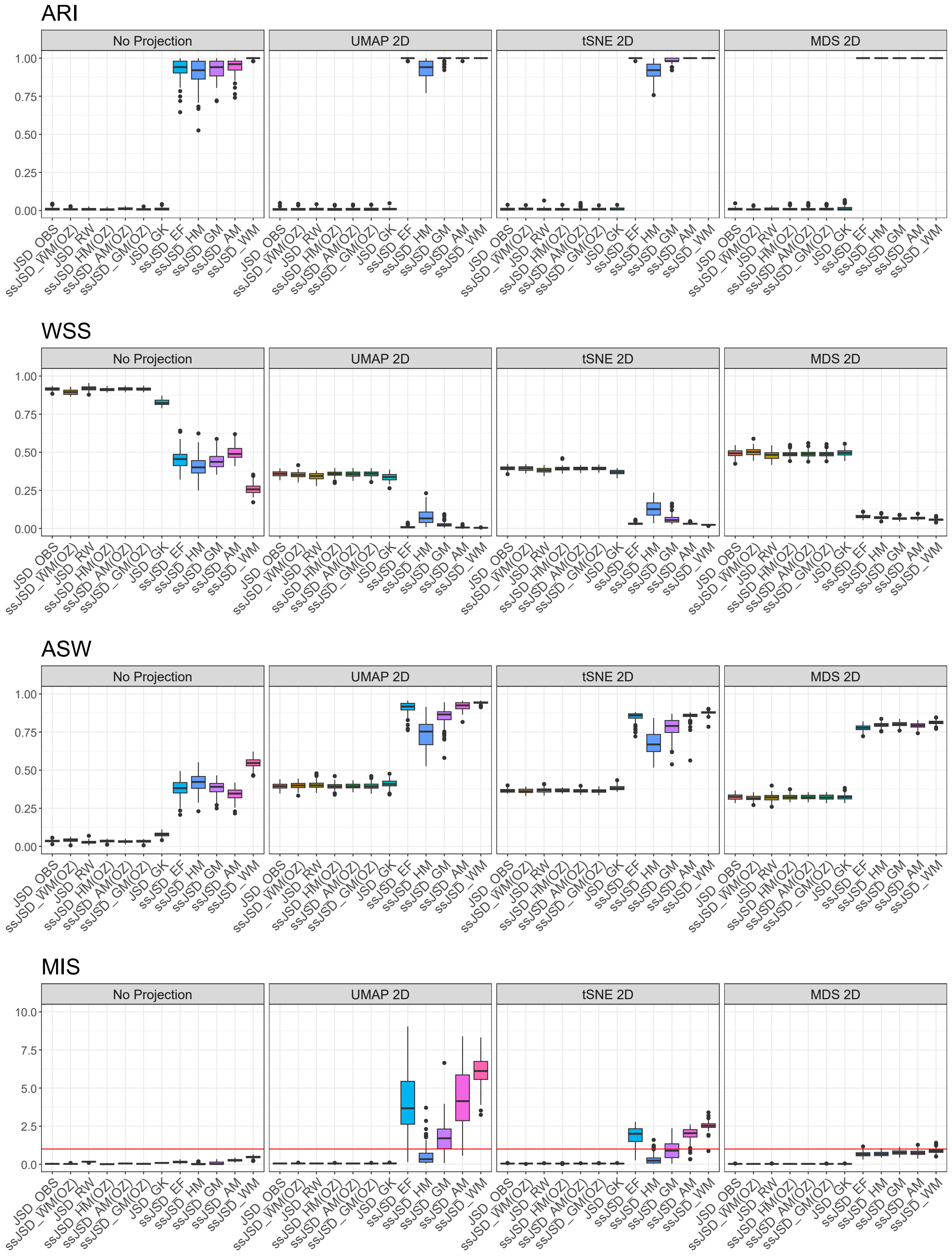


Figure S14: Comparison of the performance of 12 methods based on four clustering criteria—ARI, WSS, ASW, and MIS—across four data representations: the dissimilarity matrix (no projection), or three 2D projections (UMAP, tSNE, and MDS). The boxplots show the results across 100 datasets under Design 1 Sim3.

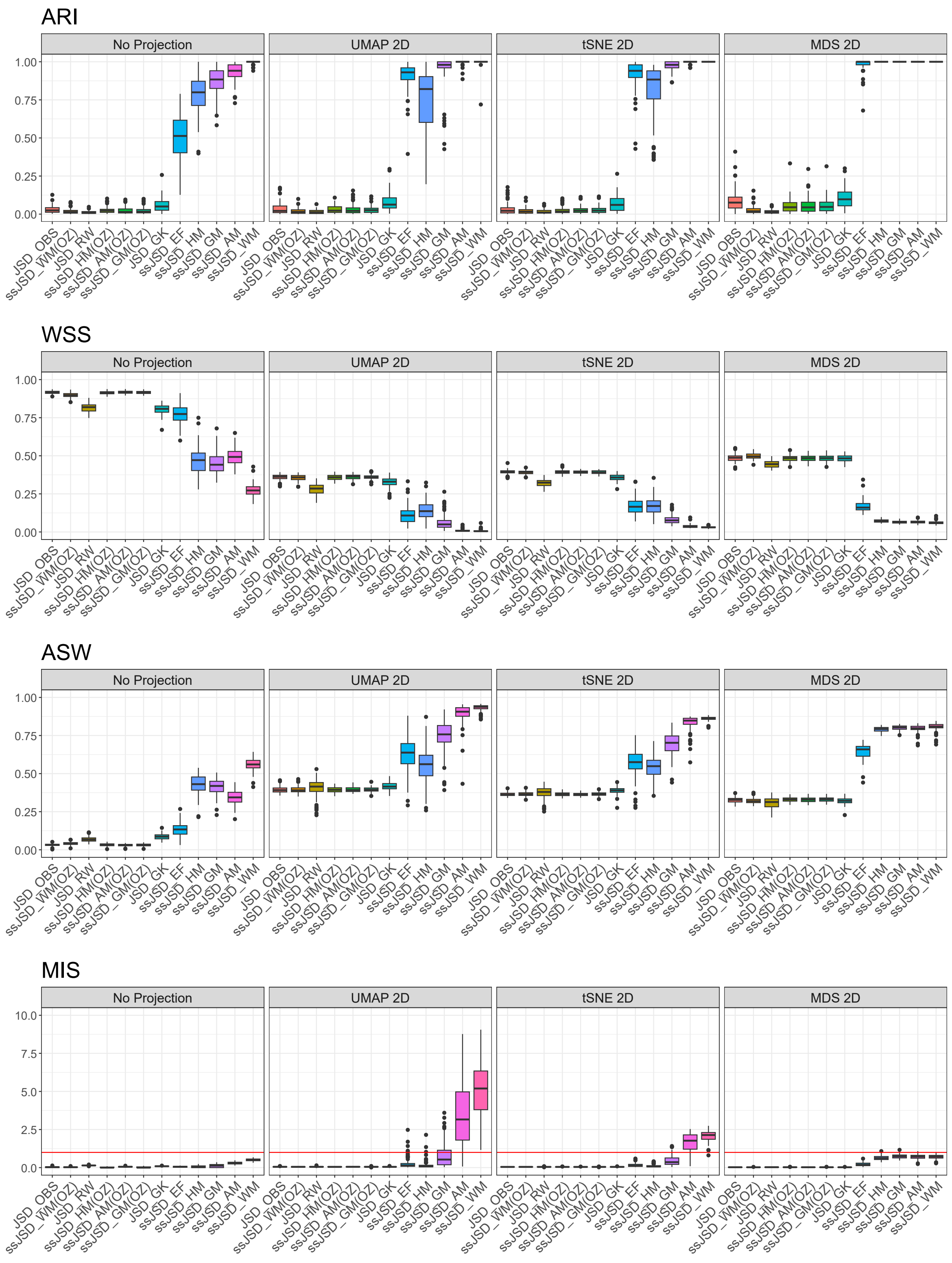


Figure S15: Comparison of the performance of 12 methods based on four clustering criteria—ARI, WSS, ASW, and MIS—across four data representations: the dissimilarity matrix (no projection), or three 2D projections (UMAP, tSNE, and MDS). The boxplots show the results across 100 datasets under Design 1 Sim4.

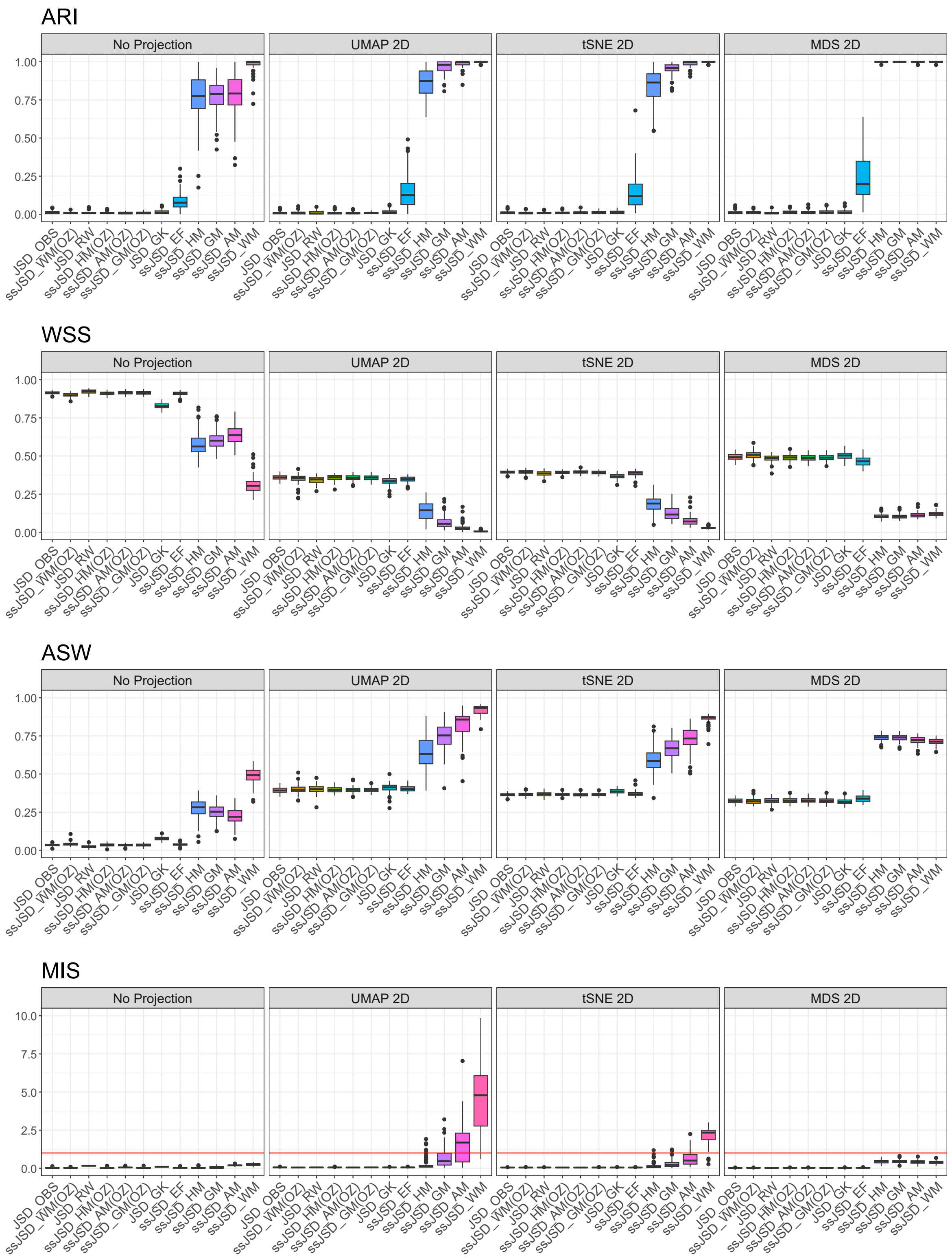


Figure S16: Comparison of the performance of 12 methods based on four clustering criteria—ARI, WSS, ASW, and MIS—across four data representations: the dissimilarity matrix (no projection), or three 2D projections (UMAP, tSNE, and MDS). The boxplots show the results across 100 datasets under Design 1 Sim5.

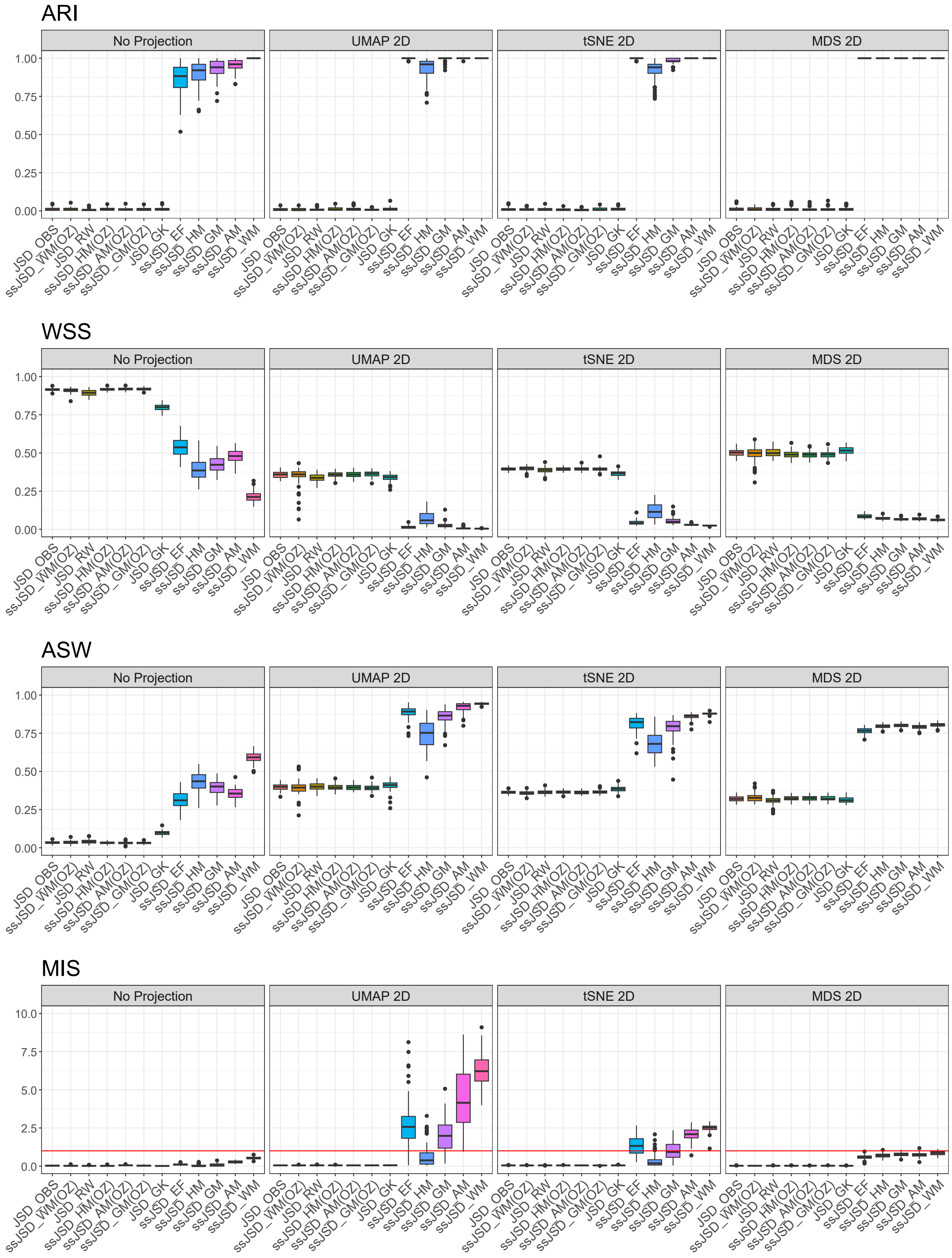


Figure S17: Comparison of the performance of 12 methods based on four clustering criteria—ARI, WSS, ASW, and MIS—across four data representations: the dissimilarity matrix (no projection), or three 2D projections (UMAP, tSNE, and MDS). The boxplots show the results across 100 datasets under Design 1 Sim6.

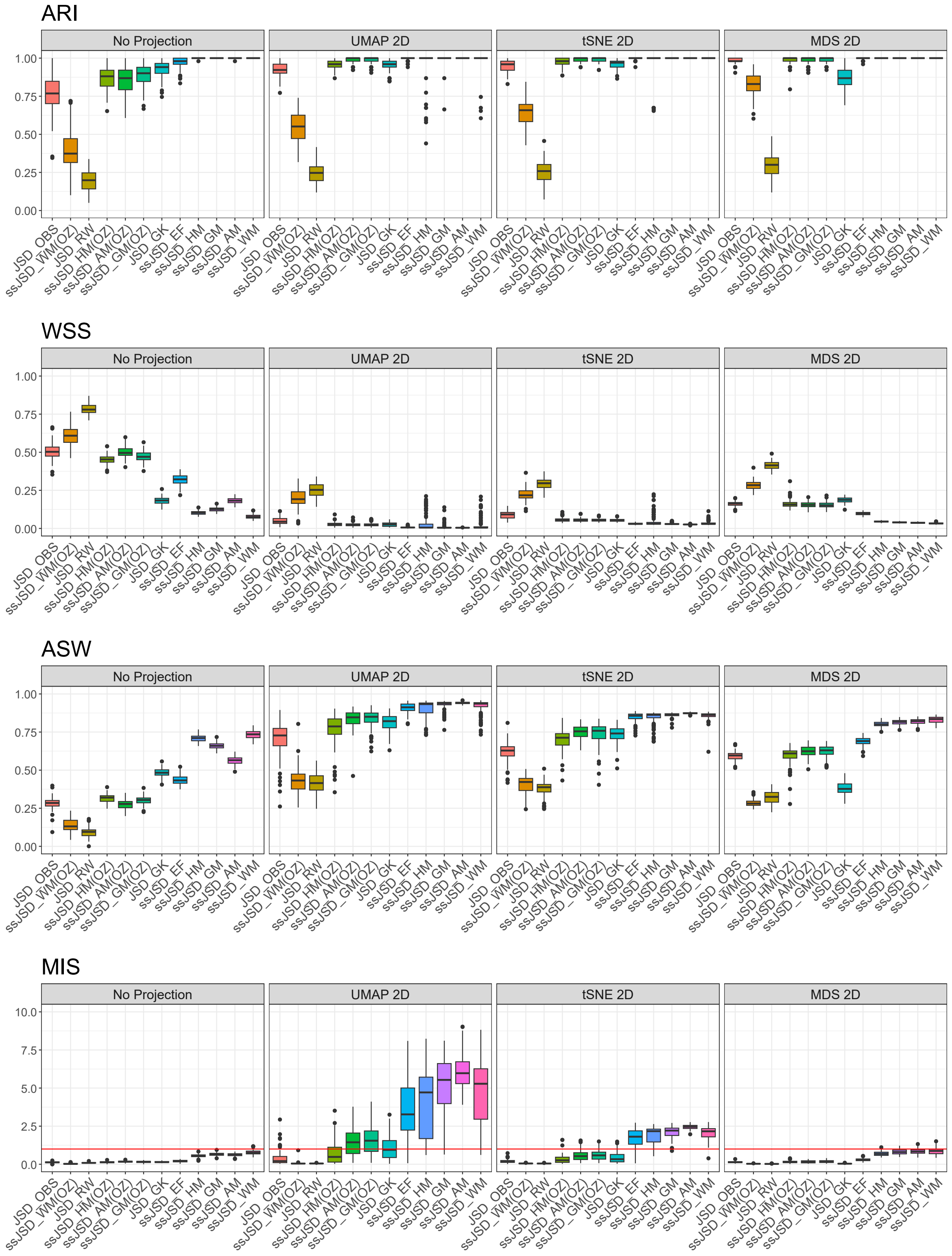


Figure S18: Comparison of the performance of 12 methods based on four clustering criteria—ARI, WSS, ASW, and MIS—across four data representations: the dissimilarity matrix (no projection), or three 2D projections (UMAP, tSNE, and MDS). The boxplots show the results across 100 datasets under Design 2 Sim1.

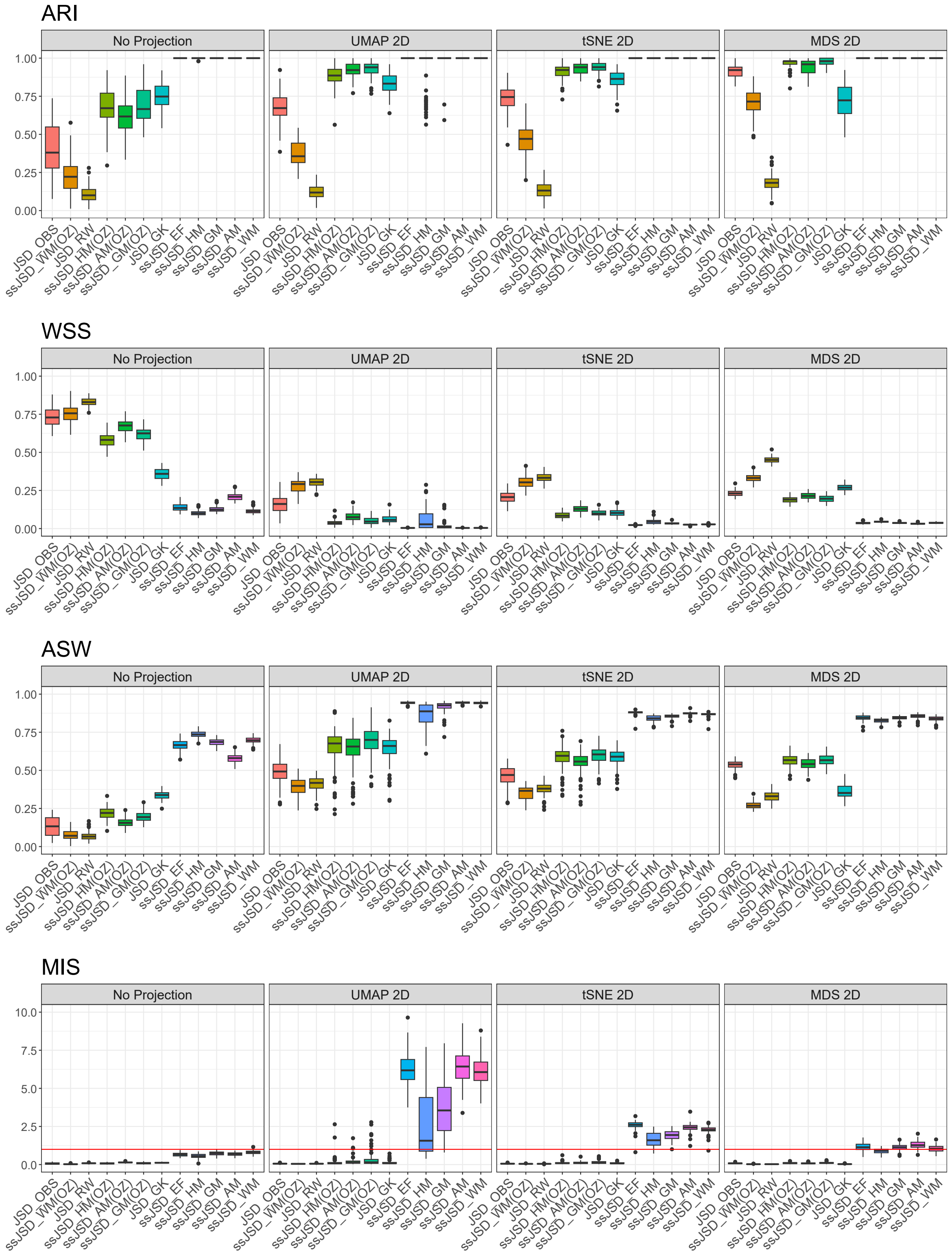


Figure S19: Comparison of the performance of 12 methods based on four clustering criteria—ARI, WSS, ASW, and MIS—across four data representations: the dissimilarity matrix (no projection), or three 2D projections (UMAP, tSNE, and MDS). The boxplots show the results across 100 datasets under Design 2 Sim2.

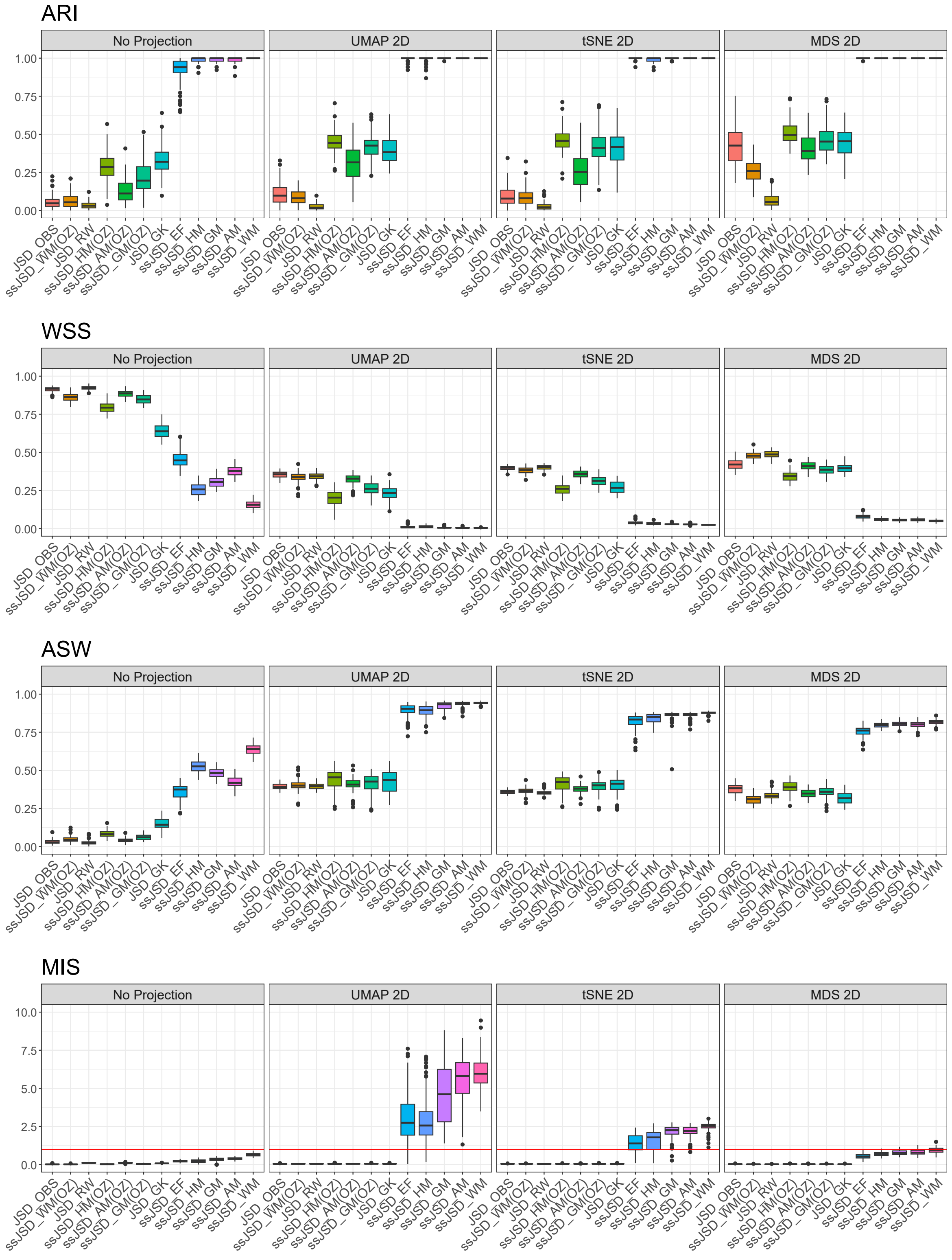


Figure S20: Comparison of the performance of 12 methods based on four clustering criteria—ARI, WSS, ASW, and MIS—across four data representations: the dissimilarity matrix (no projection), or three 2D projections (UMAP, tSNE, and MDS). The boxplots show the results across 100 datasets under Design 2 Sim3.

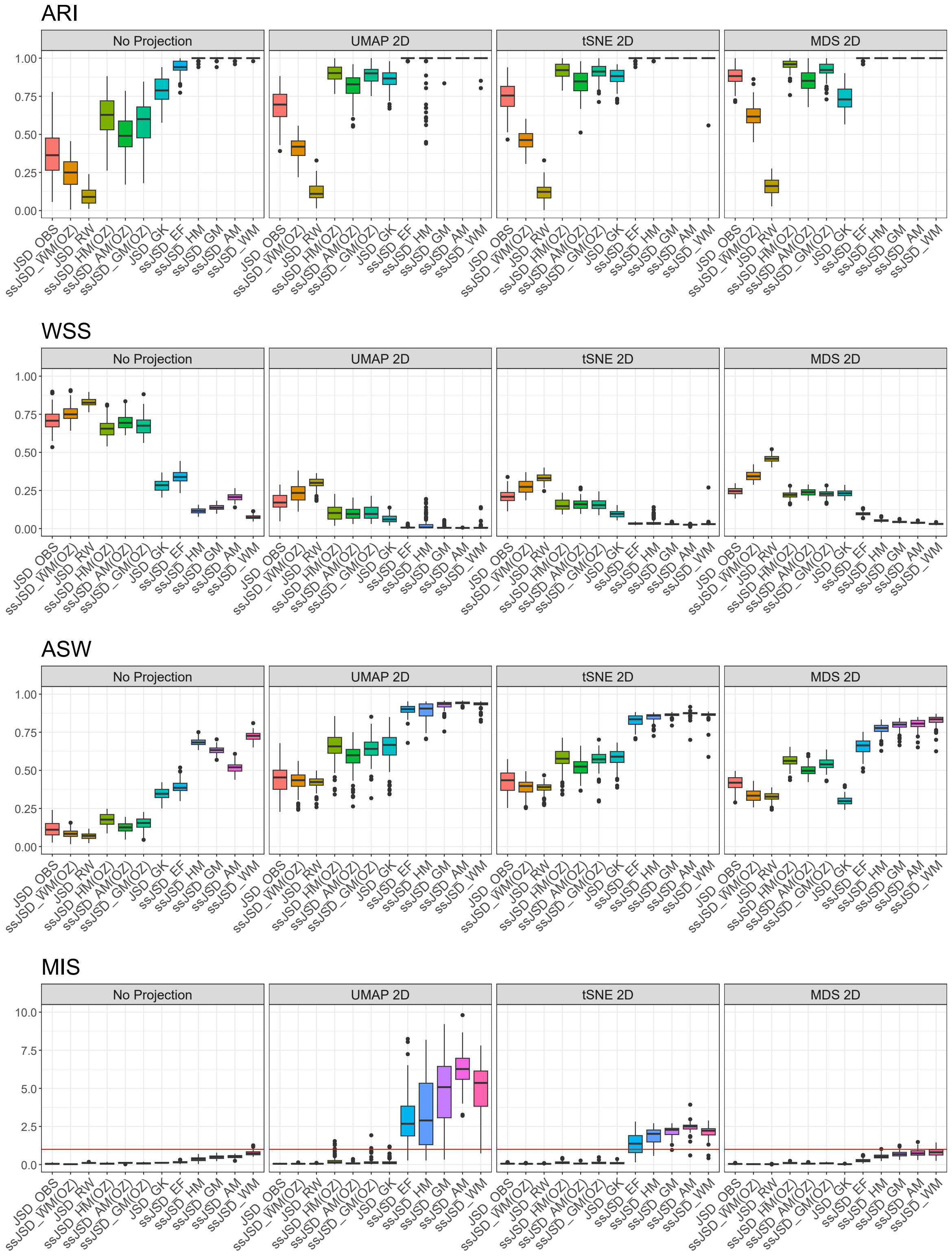


Figure S21: Comparison of the performance of 12 methods based on four clustering criteria—ARI, WSS, ASW, and MIS—across four data representations: the dissimilarity matrix (no projection), or three 2D projections (UMAP, tSNE, and MDS). The boxplots show the results across 100 datasets under Design 2 Sim4.

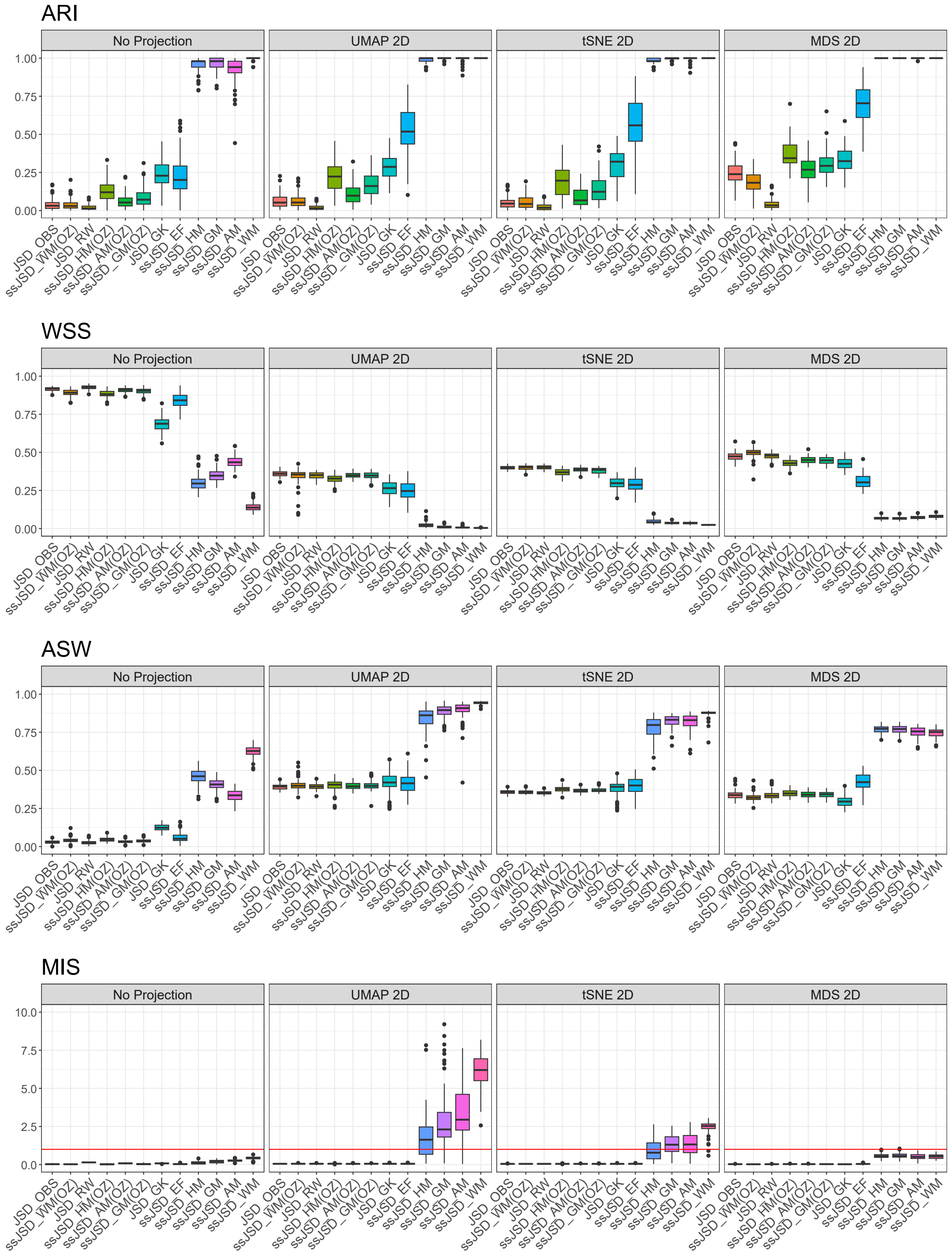


Figure S22: Comparison of the performance of 12 methods based on four clustering criteria—ARI, WSS, ASW, and MIS—across four data representations: the dissimilarity matrix (no projection), or three 2D projections (UMAP, tSNE, and MDS). The boxplots show the results across 100 datasets under Design 2 Sim5.

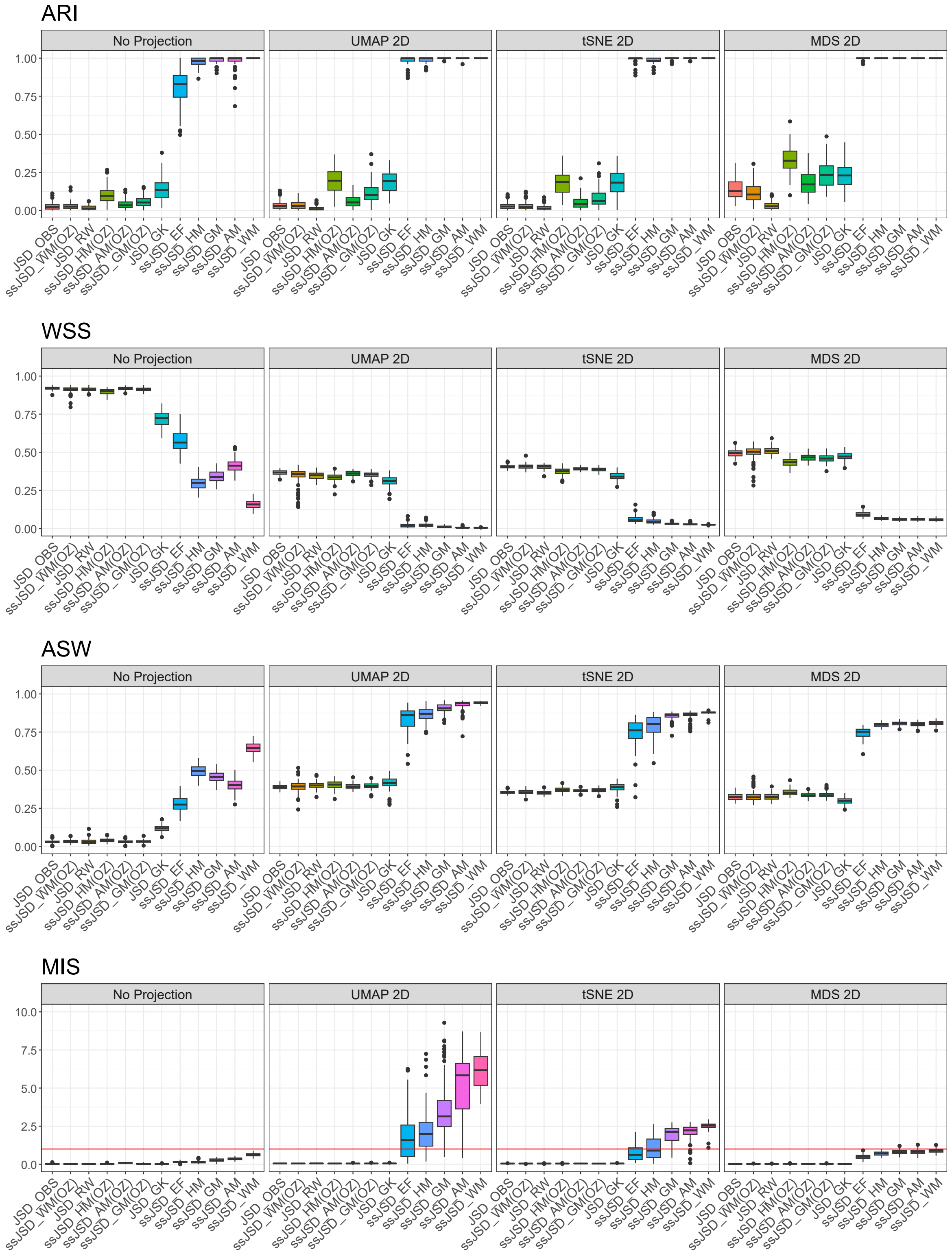


Figure S23: Comparison of the performance of 12 methods based on four clustering criteria—ARI, WSS, ASW, and MIS—across four data representations: the dissimilarity matrix (no projection), or three 2D projections (UMAP, tSNE, and MDS). The boxplots show the results across 100 datasets under Design 2 Sim6.

### (a) Design 1

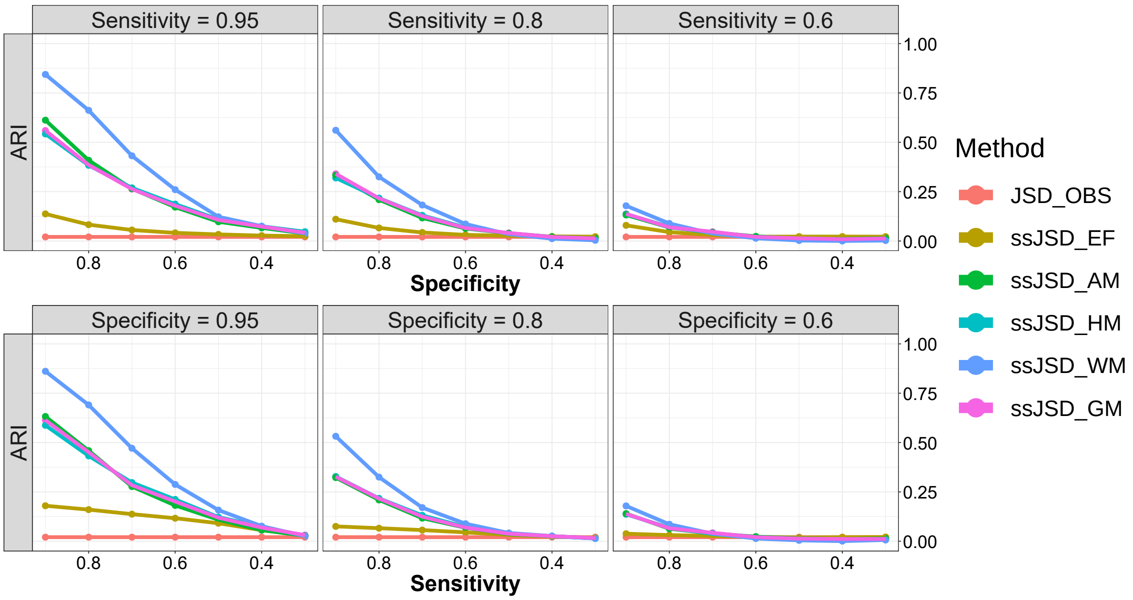


### (b) Design 2

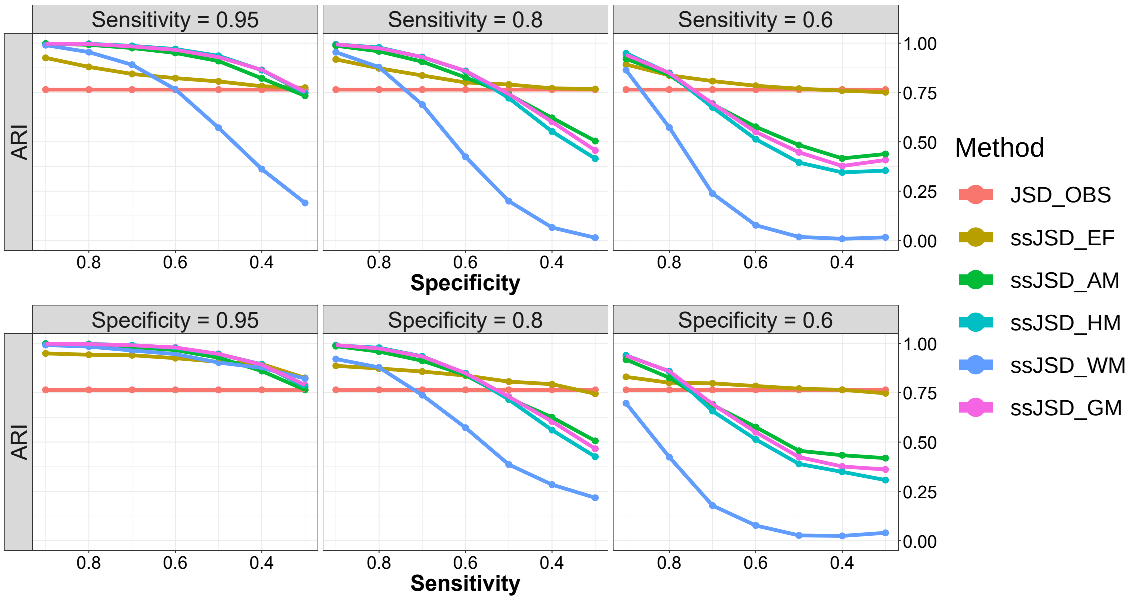


Figure S24: Effect of specificity and sensitivity on clustering performance based on ARI. (a) Design 1 Sim1 data, where each plot in the first row has a fixed sensitivity but varying specificity from 0.9 to 0.3 with a decrement of 0.1, and each plot in the second row has a fixed specificity but varying sensitivity from 0.9 to 0.3 with a decrement of 0.1; (b) Same as (a) but for Design 2 Sim1 data.

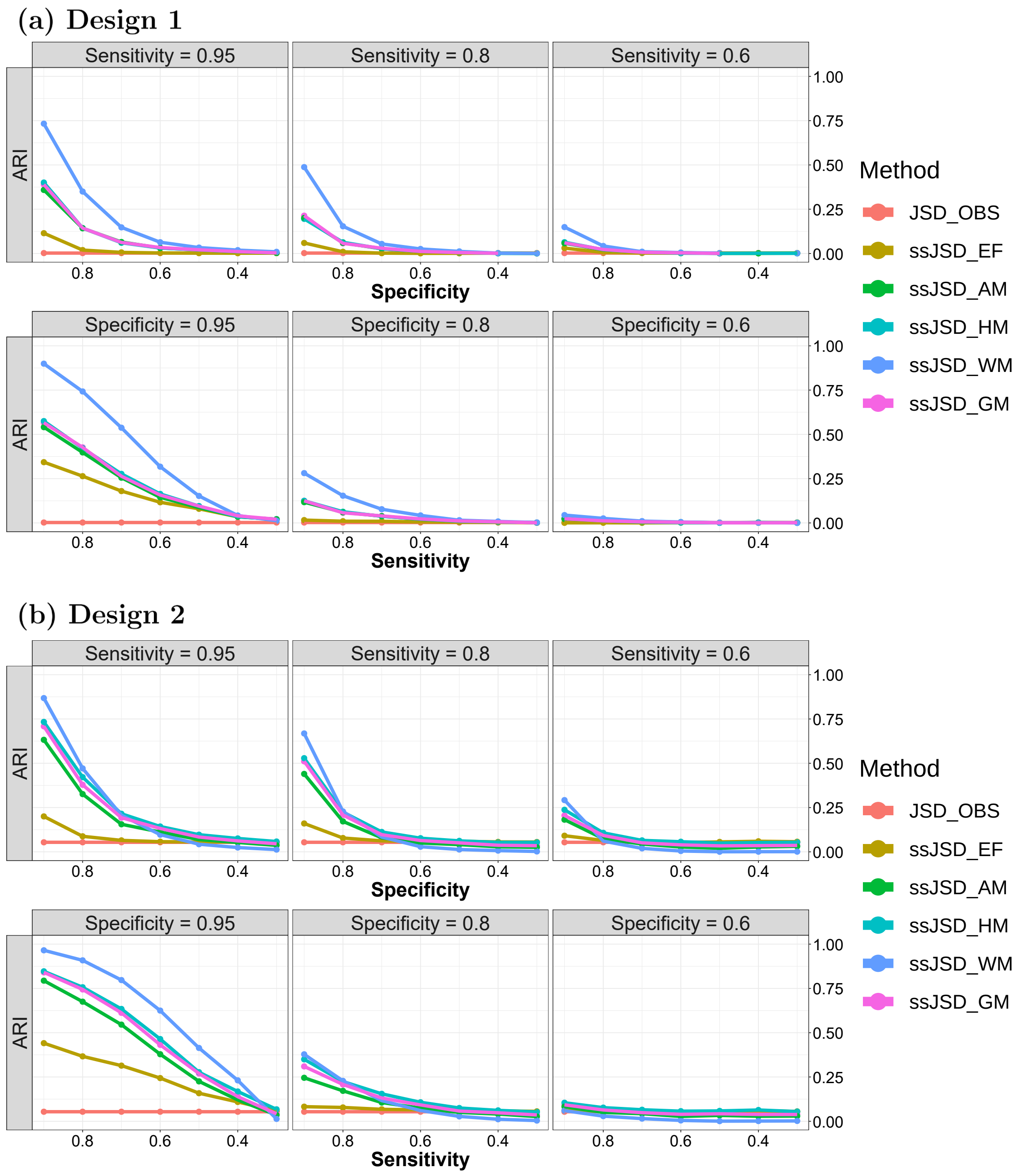


Figure S25: Effect of specificity and sensitivity on clustering performance based on ARI. (a) Design 1 Sim3 data, where each plot in the first row has a fixed sensitivity but varying specificity from 0.9 to 0.3 with a decrement of 0.1, and each plot in the second row has a fixed specificity but varying sensitivity from 0.9 to 0.3 with a decrement of 0.1; (b) Same as (a) but for Design 2 Sim3 data.

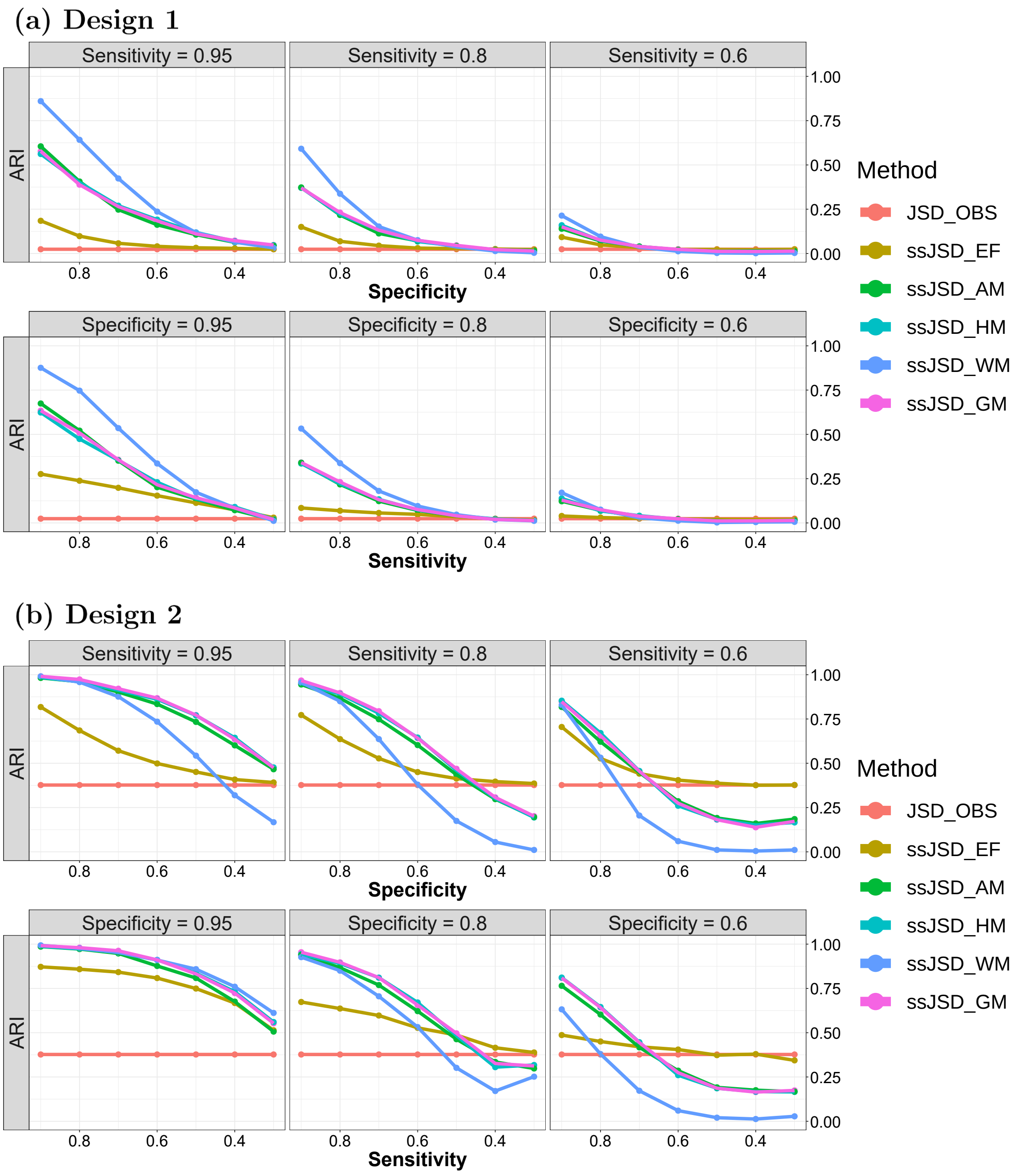


Figure S26: Effect of specificity and sensitivity on clustering performance based on ARI. (a) Design 1 Sim4 data, where each plot in the first row has a fixed sensitivity but varying specificity from 0.9 to 0.3 with a decrement of 0.1, and each plot in the second row has a fixed specificity but varying sensitivity from 0.9 to 0.3 with a decrement of 0.1; (b) Same as (a) but for Design 2 Sim4 data.

**(a) Design 1**

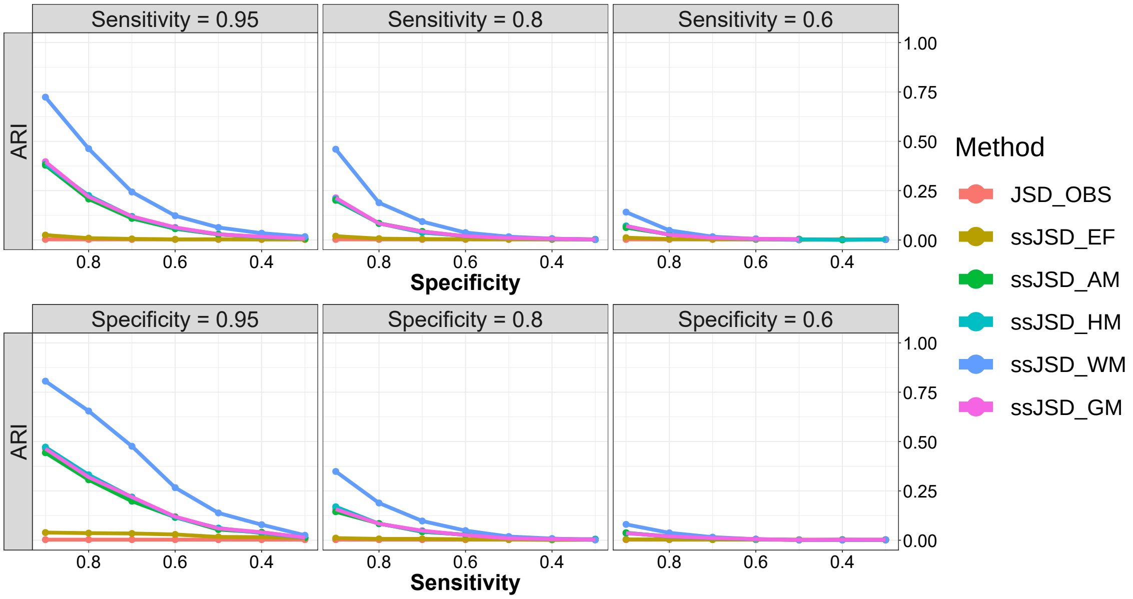


**(b) Design 2**

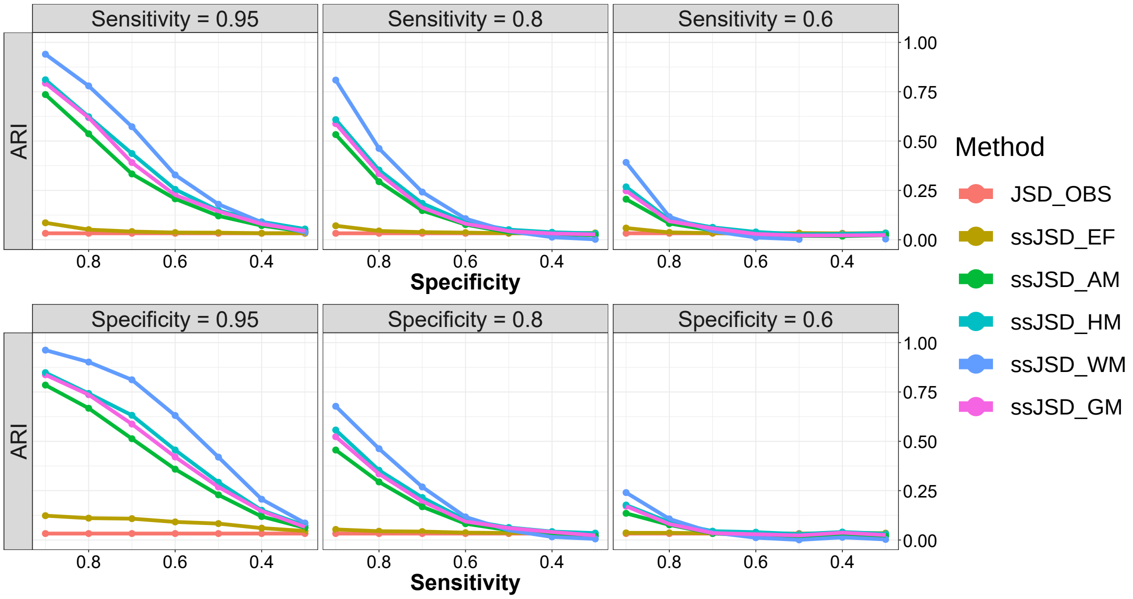


Figure S27: Effect of specificity and sensitivity on clustering performance based on ARI. (a) Design 1 Sim5 data, where each plot in the first row has a fixed sensitivity but varying specificity from 0.9 to 0.3 with a decrement of 0.1, and each plot in the second row has a fixed specificity but varying sensitivity from 0.9 to 0.3 with a decrement of 0.1; (b) Same as (a) but for Design 2 Sim5 data.

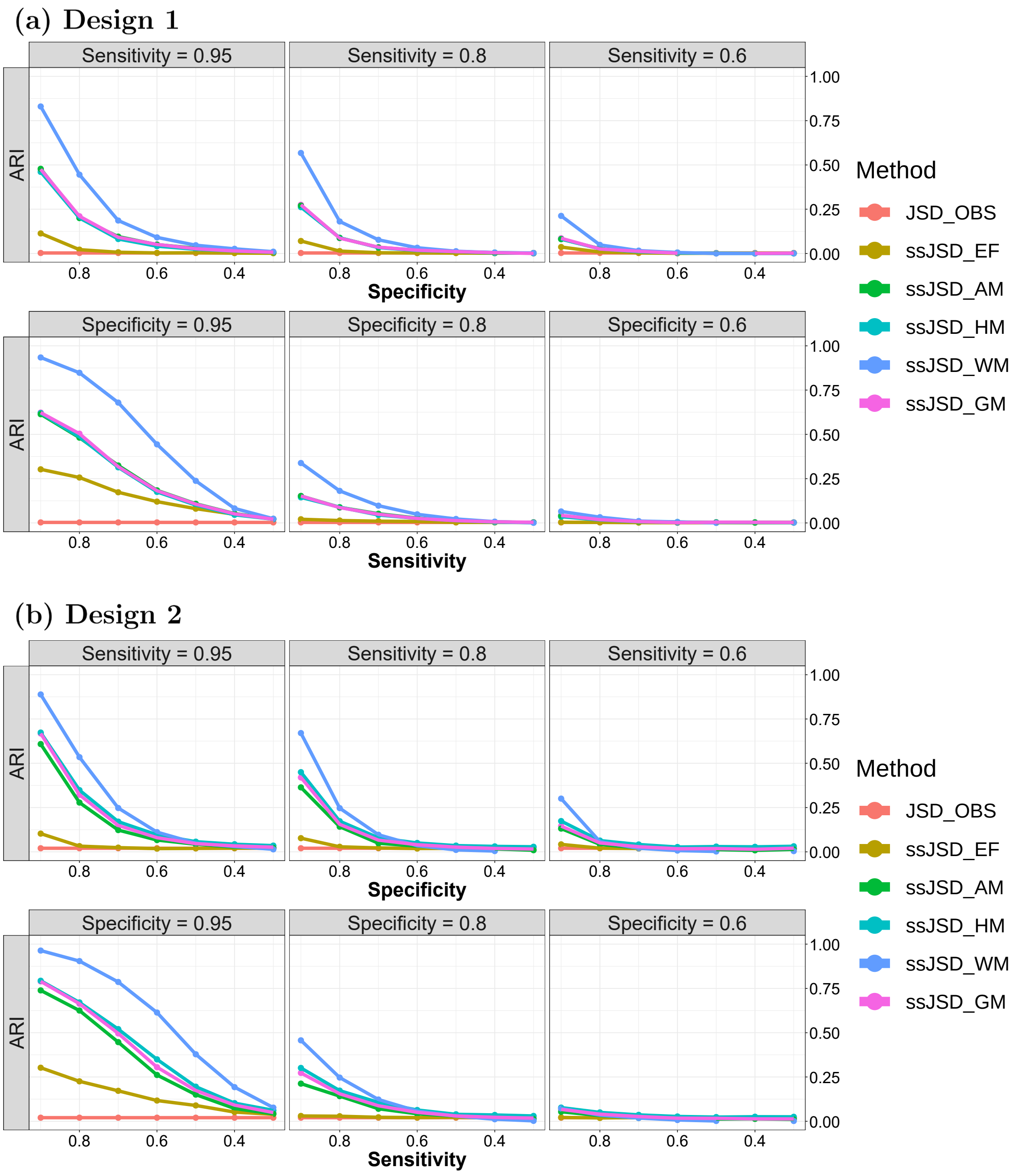


Figure S28: Effect of specificity and sensitivity on clustering performance based on ARI. (a) Design 1 Sim6 data, where each plot in the first row has a fixed sensitivity but varying specificity from 0.9 to 0.3 with a decrement of 0.1, and each plot in the second row has a fixed specificity but varying sensitivity from 0.9 to 0.3 with a decrement of 0.1; (b) Same as (a) but for Design 2 Sim6 data.

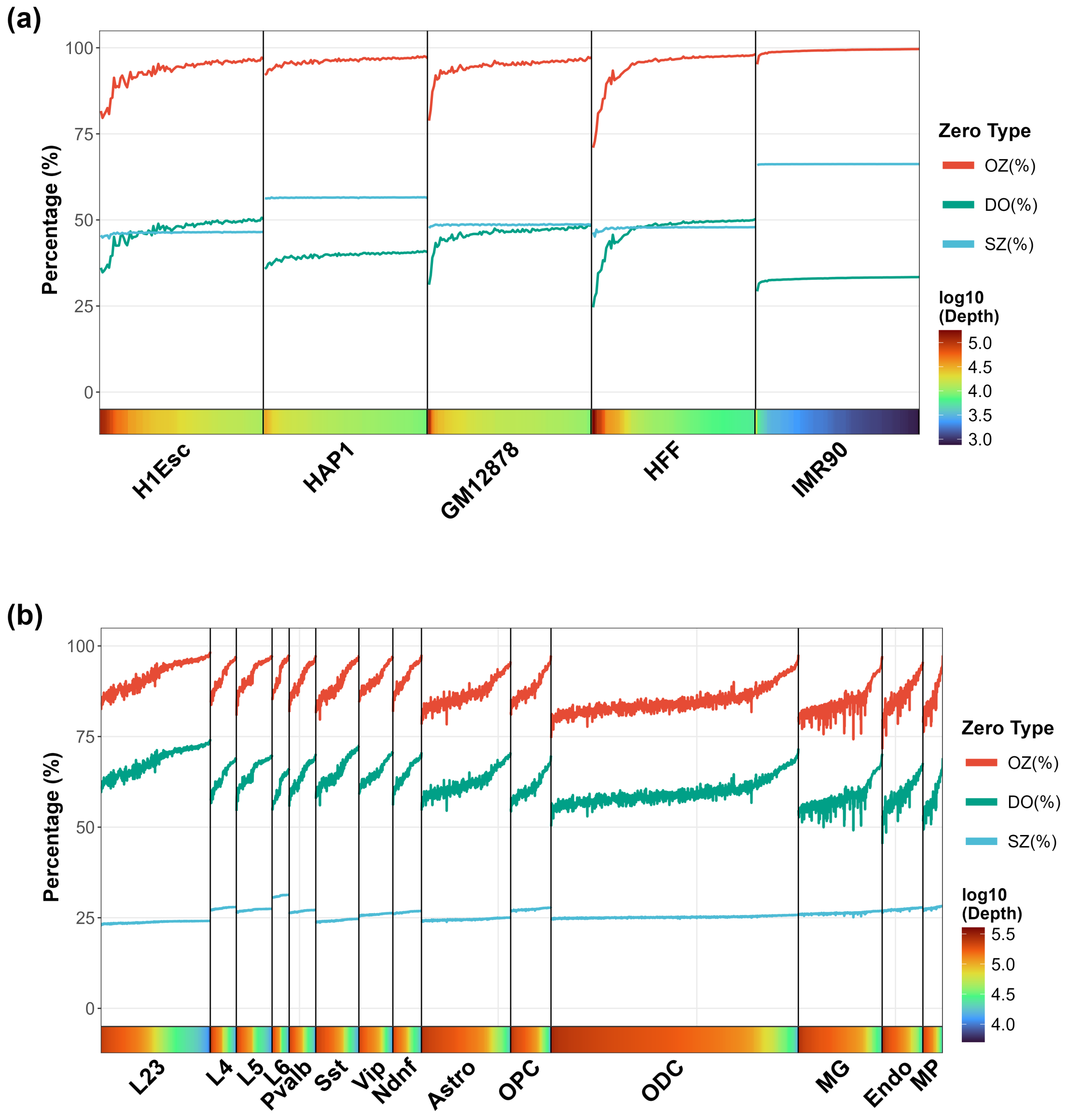


Figure S29: Results from HiCImpute on the proportions of OZs (shown for reference) being inferred to be SZs or DOs for each cell type. (a) Five human cell types data; (b) 14 cell types of the prefrontal cortex data.

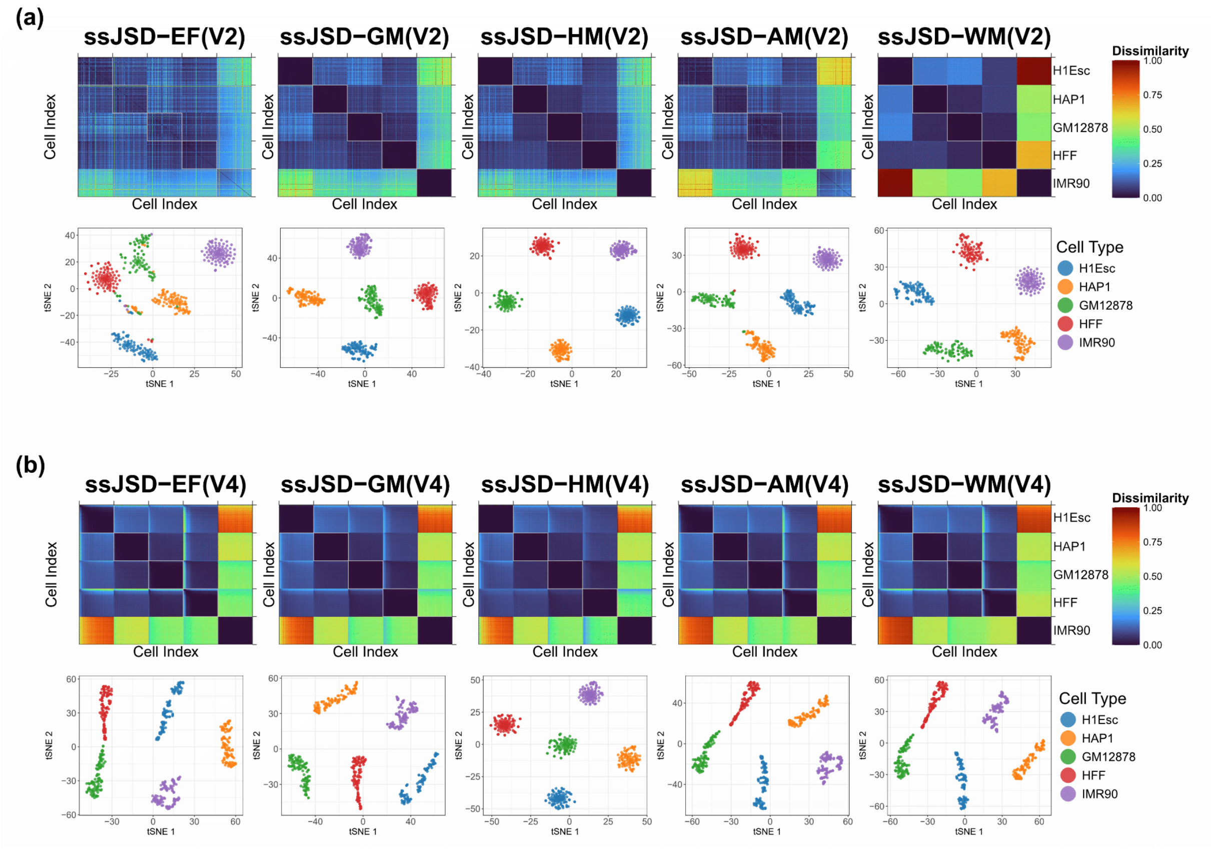


Figure S30: Results from analyzing the five human cell types data based on heatmap and 2D tSNE visualization. (a) Five ssJSD methods using V2 data (using SZ annotations only); (b) Five ssJSD methods using V4 data (SZ annotations plus imputed values for dropouts and positive counts).

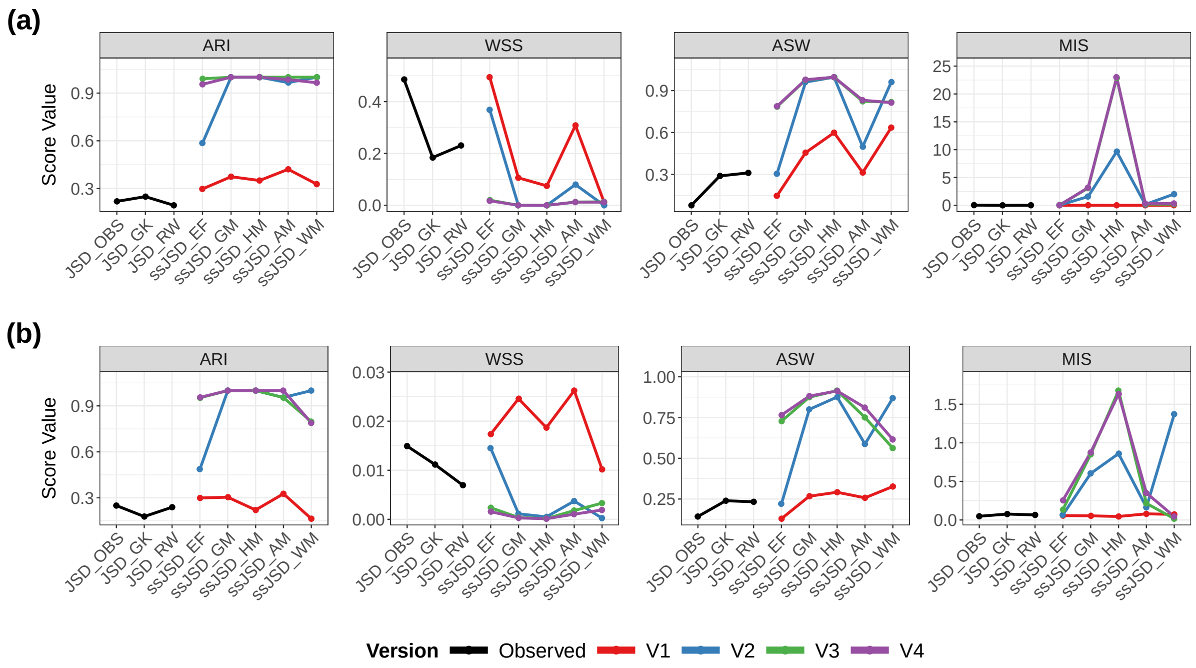


Figure S31: Performance comparison of 8 methods across four evaluation criteria (ARI, WSS, ASW, and MIS). Each line represents the utilization of different levels of imputation. (a) Results on the five human cell types data; (b) Results on the 14 human PFC data.

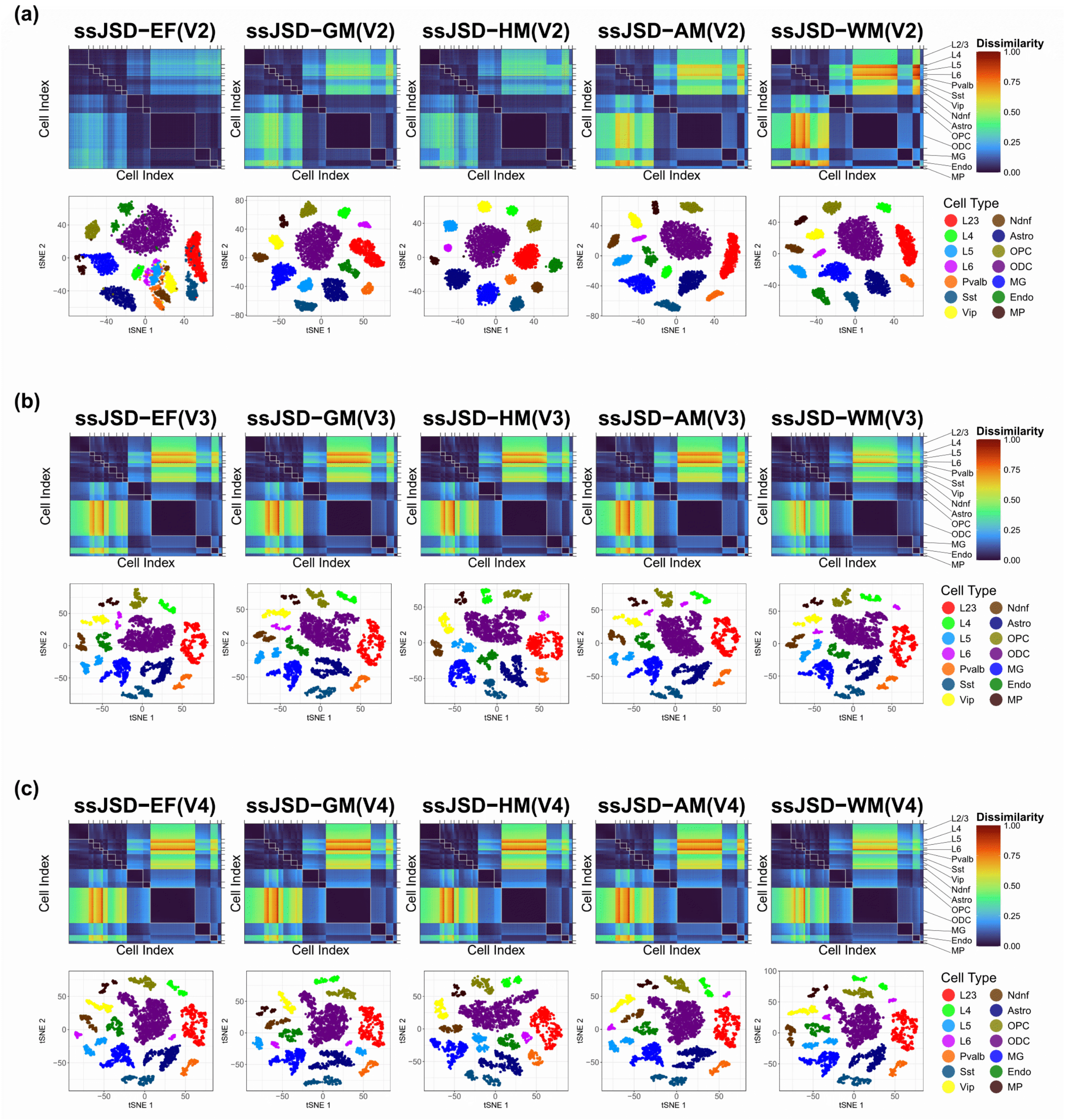


Figure S32: Results from analyzing the 14 human PFC cell types data based on heatmap and 2D tSNE visualization. (a) V2-only using SZ annotation; (b) V3-using SZ annotation plus imputed values for DOs; (c) V4-using SZ annotation as well as imputed values for DOs and positive observed counts.

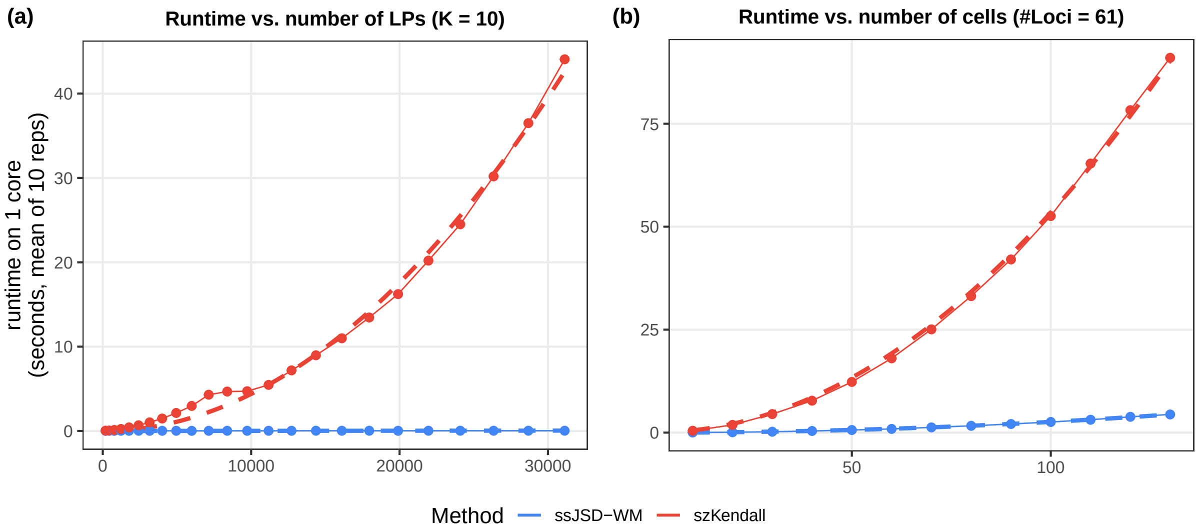


Figure S33: Runtime comparison between ssJSD and szKendall. The computational complexities for these two algorithms are in the order of $O(KL + K^2B)$ and $O(K^2L^2)$, respectively, where $K$ is the number of cells, $L$ is the number of LPs, and $B$ is the number of bands. (a) Runtime as $L$ increases, while keeping the number of cells fixed at $K = 10$, shows a quadratic curve for szKendall while linear for ssJSD, consistent with the algorithmic complexity; (b) Runtime as $K$ increases while keeping the number of loci fixed at 61 ($L = 1830$). Although both curves are quadratic, ssJSD increases much slower since its coefficient of $K^2$ is $B$, which is linear in the number of loci and thus an order of magnitude smaller than $L$, quadratic in the number of loci. The dashed lines show the fitted linear or quadratic curves consistent with the algorithm, while the solid curves are the actual computational time.
